# Activity and state-dependent modulation of salt taste behavior via pharyngeal neurons in *Drosophila melanogaster*

**DOI:** 10.1101/2022.02.01.478606

**Authors:** Shivam Kaushik, Rahul Kumar, Sachin Kumar, Srishti Sanghi, Teiichi Tanimura, Diego E. Rincon-Limas, Pinky Kain

## Abstract

Sodium present in NaCl is a fundamental nutrient required for many physiological processes. In animals, including *Drosophila,* low-salt concentrations induce attraction and high-salt concentrations evoke aversive behavior. Although the cellular basis of low and high salt taste detection has been described, the mechanisms regulating high salt consumption and salt taste modulation in animals remain elusive. Thus, we investigated the neural mechanisms behind high NaCl consumption in adult *Drosophila*, focusing on how flies adjust their acceptance of high salt based on their diet. Our findings show that long-term exposure to high salt increases the taste sensitivity of pharyngeal LSO neurons, which, in turn, enhances high salt intake. Exposing flies to a high NaCl diet for three days shows a decline in high salt aversion under starvation. Our results suggest that this modulation requires active LSO neurons and a starvation state or dopamine as genetic suppression of LSO pharyngeal neuronal activity in high NaCl-fed flies inhibit excessive salt intake. We also found that several independent taste receptor neurons and pathways are involved in such a modulation. Silencing any one of multiple LSO neuronal types inhibits excessive salt intake. Our study suggests that flies can adapt to the amount of salt ingested over several days, indicating the presence of a critical mechanism that resets the salt appetite and related neural circuits.

## Introduction

Like sugars, animals have an innate liking for low salt levels (Sodium chloride-NaCl). Sodium present in table salt (NaCl) is among the more limiting essential nutrients. Its intake must be carefully regulated to maintain ionic homeostasis, nerve signaling, fluid balance and cardiovascular activity. Animals need to ingest sodium from external food sources to carry out various physiological functions. The presence of low sodium in the body triggers specific appetite signals in the brain to drive sodium consumption. The identification of a small population of neurons in the mouse hindbrain that controls the drive to consume sodium suggests the involvement of chemosensory signals in its appetite regulation [1].

Low salt drives appetitive behavior in animals including flies while higher concentrations drive aversion [2]. Molecularly and anatomically distinct taste pathways mediate these two-opposing behavioral responses. In mice, the epithelial sodium channel (ENaC) functions as a low salt receptor [3, 4]. Interestingly, high salt does not activate its taste population in mice, but rather it co-opts other taste pathways [2] by recruiting additional pathways, including bitter and acid (sour)-sensing TRCs (Taste receptor cells). Analogous to the mammalian system, bitter GRNs (gustatory receptor neurons) respond to high salt concentrations in flies [5, 6]. In *Drosophila* larva, the *Pickpocket* (*ppk*) gene family (DEG/ENaC channels) members, *ppk11* and *ppk19,* are involved in sensing low- and high-NaCl concentrations [7] but not in adult flies [8]. Sano together with *Gr66a* has been shown to avoid high salt and is required in the terminal organs of third-instar larvae [9]

GRNs in l-type and s-type sensilla detect low- and high-NaCl concentrations in *Drosophila* labellum [8, 10-13]. *Drosophila* pharyngeal *Gr2a* taste neurons mediate aversion to high salt [14]. Additionally, *Drosophila dpr* locus (for defective proboscis extension response), a member of the Ig superfamily, is required for the aversive response to high salt [15]. TMC-1 (*trans*-membrane channels like) is a sodium channel that controls high salt aversion in *C.elegans* [16]. *Ir76b* detects low and high-NaCl concentrations [8, 13]. Moreover, a complex code for salt taste has been documented in flies [17] as well. Also, adult female *Drosophila* switches from sugar feeding to fermenting yeast to get protein and micronutrients for egg development [18, 19]. Recently, the roles of *Ir56b* in mediating appetitive behavioral responses to salt and *Ir7c* in mediating normal avoidance of high monovalent salt concentrations were reported [20, 21]. Indeed, bit was found that both *Ir56b* and *Ir7c* act with co-receptors *Ir252/Ir76b* [20, 21]. Others suggested role of bitter taste neurons and a class of glutamatergic ‘‘high salt’’ neurons expressing *pickpocket23* (*ppk23*) in driving high salt avoidance [6, 17, 22].

Although the cellular basis of low and high salt taste detection has been extensively studied, the mechanisms regulating high salt consumption and salt taste modulation in animals remain largely elusive. In a recent study, functional and behavioral experiments revealed the role of different subsets of pharyngeal neurons in governing avoidance responses to taste stimuli including high salt [23]. Other studies highlighted inhibitory mechanism for suppressing high salt intake [24] and probed whether IRs function for avoidance of high sodium and showed the role of *Ir60b* in pharyngeal GRNs along with co-receptors *Ir25a* and *Ir76b* in limiting salt consumption [25]. To understand how pre-exposure to high NaCl diet modulates subsequent taste behavior, we looked into the molecular mechanisms of a high NaCl consumption and its impact on the feeding behavior using adult *Drosophila* as a model system. Our study suggests a neural mechanism in which flies modify their acceptance of high salt as a function of diet, where a long-term high-salt exposure increases taste sensitivity of pharyngeal labral sense organ (LSO) neurons and enhances high salt intake. We also found that wild-type flies are attracted to low NaCl levels and show aversion towards higher concentrations. However, exposing these flies to a high NaCl diet for three days modifies their feeding preference to high salt levels, whereas high NaCl-fed flies show a decline in high salt aversion under starvation. Strikingly, genetic suppression of LSO pharyngeal neuronal activity in high NaCl-fed flies inhibit excessive salt intake. We found that this modulation requires active LSO neurons and a starvation state or dopamine. Our results also suggest multiple independent taste receptor neurons and pathways are involved in such a modulation as silencing any one of the multiple LSO neuronal types inhibit excessive salt intake. Our data support the idea that high dietary salt modulates and reshapes salt and other taste curves in flies. In conclusion, our study suggest that flies can adapt to the amount of salt ingested over several days, indicating the presence of a critical mechanism to reset the salt appetite and related neural circuits.

## Results

### Wildtype flies show aversion to high NaCl concentrations

To understand how flies, respond to varying concentrations of NaCl, we first tested wildtype (*CsBz*) flies for their feeding preferences in a feeding assay (**Figure 1A**). To examine the salt feeding behavior in feeding assays, batches of 20 flies per plate (10 males + 10 females) were presented with a choice between water and varying NaCl concentrations. In our two-hour feeding assay, flies were allowed to choose between these options in an unbiased manner in the dark. Blue spots in the feeding plate consisted of just water and agar. NaCl added with agar and water was presented as red spots (**Figure 1A**). % flies showing a preference for NaCl (eating red) were scored based on their abdomen color. We found that wild-type flies show highest attraction to 10, 50 and 100mM NaCl concentration exhibiting highest feeding preference at these concentrations (**Figure 1B**). But with the increase in salt concentration, flies show reduced feeding preferences toward high NaCl concentrations (200mM, and 500mM) (**Figure 1B**). Our dye switching experiment (NaCl presented in blue spots) also showed the same feeding preference at 10, 50 and 100mM NaCl (**Supplementary Figure 1A**).

**FIGURE 1.**
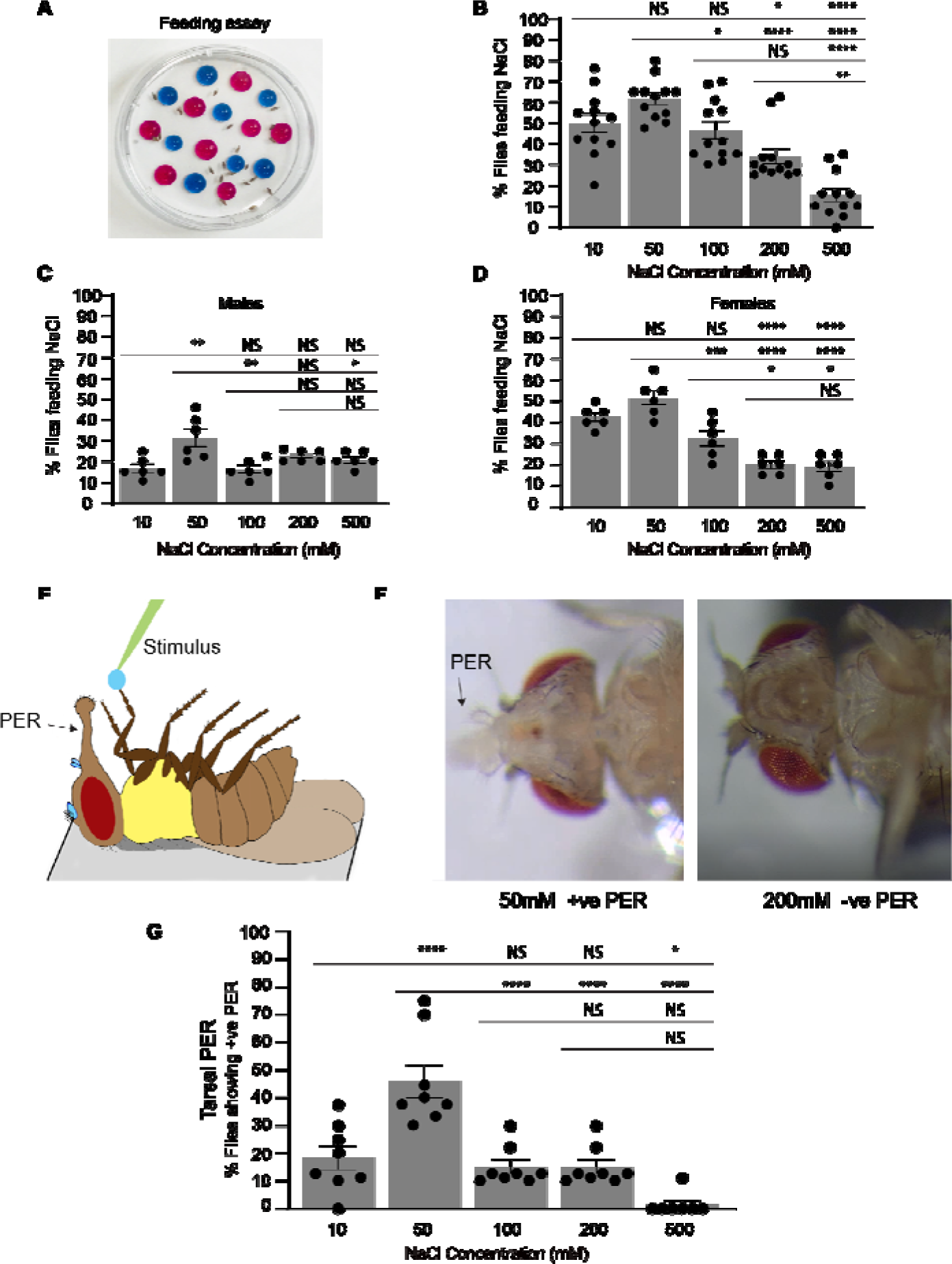
Flies show attraction to low concentrations of salt (NaCl) and aversive behavior towards high salt concentrations. (**A**) Image of feeding assay plate. Feeding assay plate showing red spots having salt and agar. Blue spots correspond to control dots containing water and agar. (**B**) Dose response profile showing mean NaCl feeding preference of wild type flies (*CsBz*) at the indicated concentrations of NaCl in the feeding assay after 24 h of starvation. For each bar, n=12 trails of ∼20 flies each for each concentration (10 males and 10 females). (**C** and **D**) Dose response profile showing % mean NaCl feeding preference of starved wild type mated male (**C**) and mated female flies (**D**) at different concentrations of NaCl in the feeding assay. For each graph, each bar, n=6 trails of 20 flies each for each concentration. (**E**) Schematic of a fly showing tarsal proboscis extension reflex (PER) response in response to a positive stimulus. (**F**) Bright field images of flies showing extension of proboscis (arrow) at an attractive NaCl concentration (50mM NaCl, left) and no extension at higher concentration (200mM NaCl, right). (**G**) Dose response profile showing % mean positive PER responses of wild type flies (*CsBz*) at the indicated concentrations of NaCl in the Tarsal PER assay. For PER, n= 72 flies. Dead or immobile flies were not considered or used for the assay. % Flies showing positive PER was calculated for 8 batches of flies. Mix of males and females were used in each case. Statistical analysis was performed using ANOVA Tukey’s multiple comparison test for obtaining P values: *p < 0.05, **p < 0.005 and ***p < 0.0005. For all graphs, error bars=SEM and NS is not significant.

We observed female flies show slightly high preference for NaCl, when mated males and females were tested separately in the feeding assays. (**Figure 1C, D**) suggesting special requirement of sodium ions to invest in the progeny in case of female flies [26]. In both male and female flies (**Figure 1C, D**), the highest feeding preference was observed at 50mM NaCl concentration. We found 50mM concentration showed the most significant differences when compared with other NaCl concentrations (**Figure 1B, C, D**).

We then used an independent assay to assess if flies prefer lower concentrations of NaCl and show aversive responses at higher NaCl concentrations. We stimulated the tarsal taste hairs of wildtype (*CsBz*) flies to perform the tarsal PER (Proboscis Extension Reflex assay, **Figures 1E, F**) assay with a series of NaCl concentrations ranging from 10-500mM (same concentrations used in the feeding assays). As observed in our feeding results, we observed the highest proboscis extension response (∼ 47%) in wild-type flies (mix of males and females) at 50mM NaCl concentration (**Figure 1G, Supplementary video 1**) and most significant differences with other NaCl concentrations. During this assay flies were not allowed to drink the taste solution. Also, while performing the assay flies were monitored on the screen to ensure that they were not drinking the solution. We observed 15-20% positive PER responses in case of tarsal PER (**Figure 1G**) at 10, 100 and 200 mM (**Supplementary video 2**) concentrations. Less than 10% responses were recorded at the highest concentration (500mM) tested (**Figure 1G**) suggesting reduced attraction at high concentrations. Snapshots of tarsal PER responses at 50mM (positive PER response showing extended proboscis, left image in **Figure 1F**, **Supplementary video 1**) and 200mM NaCl concentrations (negative PER-right image showing no proboscis extension in **Figure 1F and Supplementary video 2**). These results suggest that flies show attractive behavior to low concentration (50mM) of NaCl and avoid NaCl as the concentration increases. In another set of PER experiments, we also tested sugar at the beginning and at the end of the experiments to make sure flies were not losing sensitivity after testing various salt concentrations. We plotted responses of 85-95% flies (positive PER) that responded to sucrose before and after presenting the NaCl (**Supplementary Figures 1B).**

### High NaCl fed flies show a decline in high salt aversion under starvation

To determine if prior exposure to high NaCl alters the feeding behavior for the preferred level of salt, wild-type flies were fed continuously for 3 days on high salt diet (200 mM NaCl mixed with standard fly food; **Figure 2A**,). We decided a time period of 3 days based on our results obtained with 1- and 2-day feeding (**Supplementary Figures 2A, B**). Following starvation period of 24 h, flies were tested in a feeding assay for their feeding preference for different concentrations of NaCl (**Figures 2A, B**). In these experiments, flies kept on standard fly media for 3 days were treated as control flies (**Figures 2A, B**). We observed a shift in feeding preferences, flies preferring higher concentrations of NaCl when kept on high salt media. These flies maintained their preferences for high NaCl concentrations and showed increased feeding preference for 100 and 200mMm NaCl concentrations (black bars in **Figure 2B**). No difference in feeding was observed at lower concentrations (10 and 50mM, **Figure 2B**). As we found reduced aversion towards high NaCl when wildtype (*CsBz*) flies were pre-exposed to a high NaCl diet in our feeding assays, later we confirmed feeding results by performing spectrophotometry on flies (**Figures 2C, D**) to determine food intake. In our food intake spectrophotometry experiments, the absorbance values showed similar results when two different wildtype flies (*CsBz* and *w^1118^*) were tested (**Figures 2C, D**). Even in these experiments, interestingly we found flies pre-exposed to high NaCl showed reduced repulsion towards high NaCl when compared to standard fly media flies.

**FIGURE 2.**
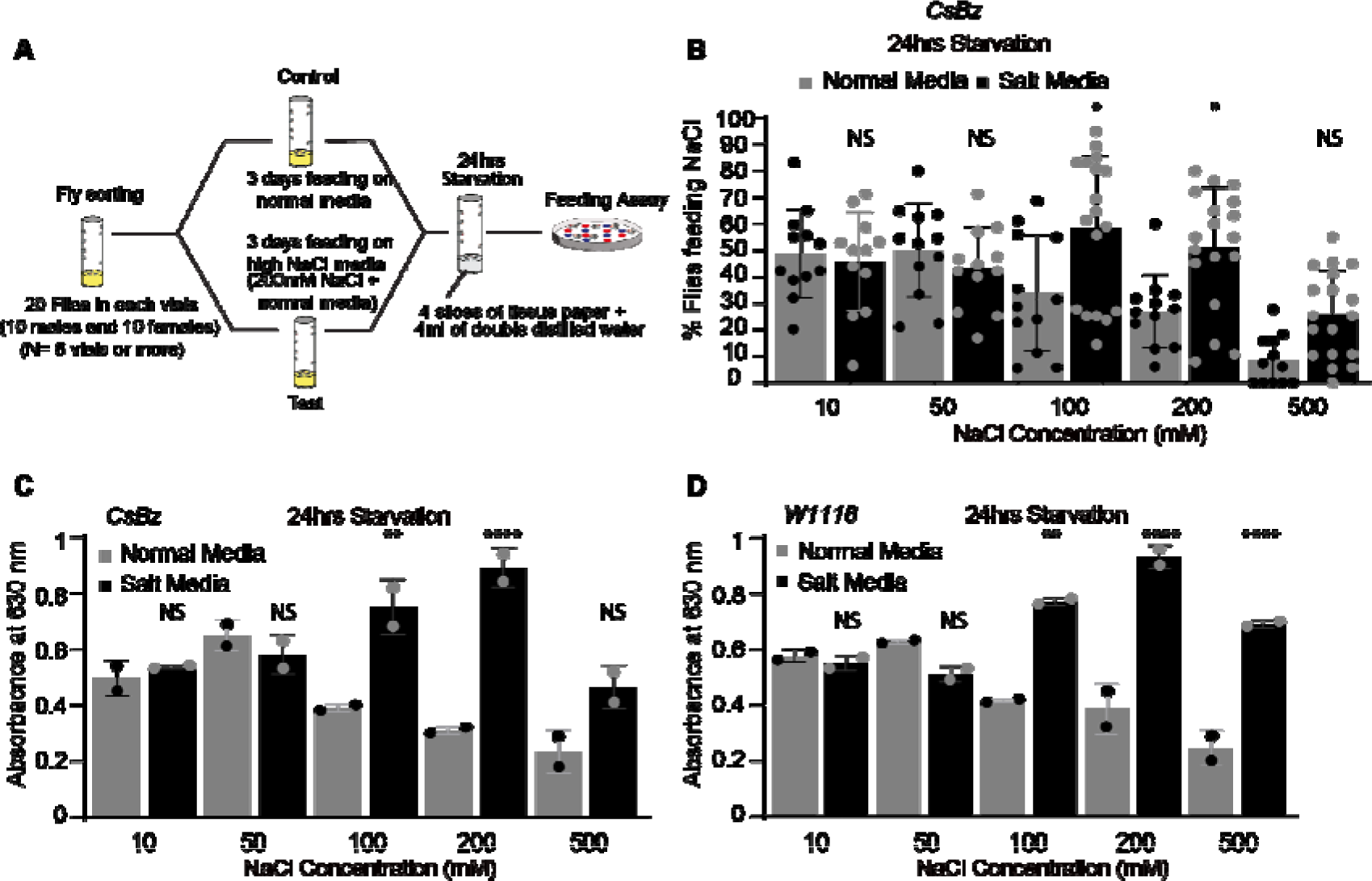
High NaCl fed flies show decline in high salt aversion under starvation. (**A**) Schematic of feeding paradigm used to pre-expose wild type flies to high NaCl condition. (**B**) Dose response profile of *CsBz* wildtype flies showing % mean NaCl feeding preferences for indicated concentrations of NaCl in two different feeding backgrounds. Black bars show preferences of flies pre-fed on high salt diet (200mM NaCl mixed with normal fly food) for 3 days compared with flies fed on normal fly media for 3 days (gray bars). After 24hrs of starvation, flies were tested to check their preferences for the indicated concentrations of NaCl. For each bar, n=12-18 trails of 20 flies each (10 males and 10 females). (**C** and **D**) Mean absorbance values for *CsBz* and *w^1118^* flies kept on normal media and high salt media. n= 2 sets each concentration, 60 flies each set. For all graphs, error bars=SEM. Statistical analysis was performed using ANOVA Tukey’s multiple comparison test for obtaining P values: *p < 0.05, **p < 0.005 and ***p < 0.0005. For all graphs, error bars=SEM and NS is not significant.

We also tested if change in the internal state was responsible for modulating the salt taste behavior in the high salt-fed flies. We tested wildtype (*CsBz*) flies kept on high NaCl media for 3 days and on standard media conditions (control condition) without any starvation (**Supplementary Figure 3A**). In the fed state, we observed no significant differences in feeding preference between standard media and high salt-fed flies (**Supplementary Figure 3A**). Even under fed conditions, flies showed highest attraction to salt at 50mM concentration (**Supplementary Figure 3A** grey bars). Under fed condition, the overall feeding preferences were reduced compared to starved flies (**Supplementary Figure 3A** grey bars and **Figure 2B**). Our data suggest that internal state plays a role in modulating salt taste behavior and guide the fly feeding decisions under starved or fed conditions.

### High NaCl fed flies show no alteration in consumption of potassium under starvation

Sodium works alongside potassium in animals to conduct electrical impulses, muscle contraction, and maintain many other important physiological functions. Animals, including humans, require daily need of potassium to support these key processes. Next, we tested if flies feeding on high NaCl media also need high potassium. We tested flies feeding on standard and high NaCl diet to determine their feeding preferences for different concentrations of KCl (potassium chloride; **Supplementary Figure 3B**). We found that high NaCl-fed flies did not consume higher KCl at any tested concentration and no significant differences in the feeding preference of flies were observed on normal media and high NaCl media conditions (**Supplementary Figure 3B**). Our data suggest that pre-exposing flies to high-NaCl diet does not affect KCl consumption.

### High NaCl fed flies show alteration in consumption of metabolizable sugars

To understand if pre-exposure to a high NaCl diet causes any alteration in preferences for other taste modalities, we tested sweet taste. Varying concentrations (50, 100, and 200mM) of sucrose were presented to flies and tested in our feeding assays. We didn’t observe significant differences in consumption and feeding behavior between flies fed on standard media or high NaCl media (**Supplementary Figures 4A, B**). Only when lower concentrations of sucrose were presented to flies in the feeding assay and consumption was monitored using spectrophotometry analysis, we observed increased consumption at 5, 10, and 15mM sucrose concentrations in high NaCl fed flies (**Figure 3A**). We observed no change in sucrose consumption at 1, 20, and 25 mM between test and control flies (**Figure 3A**).

**FIGURE 3.**
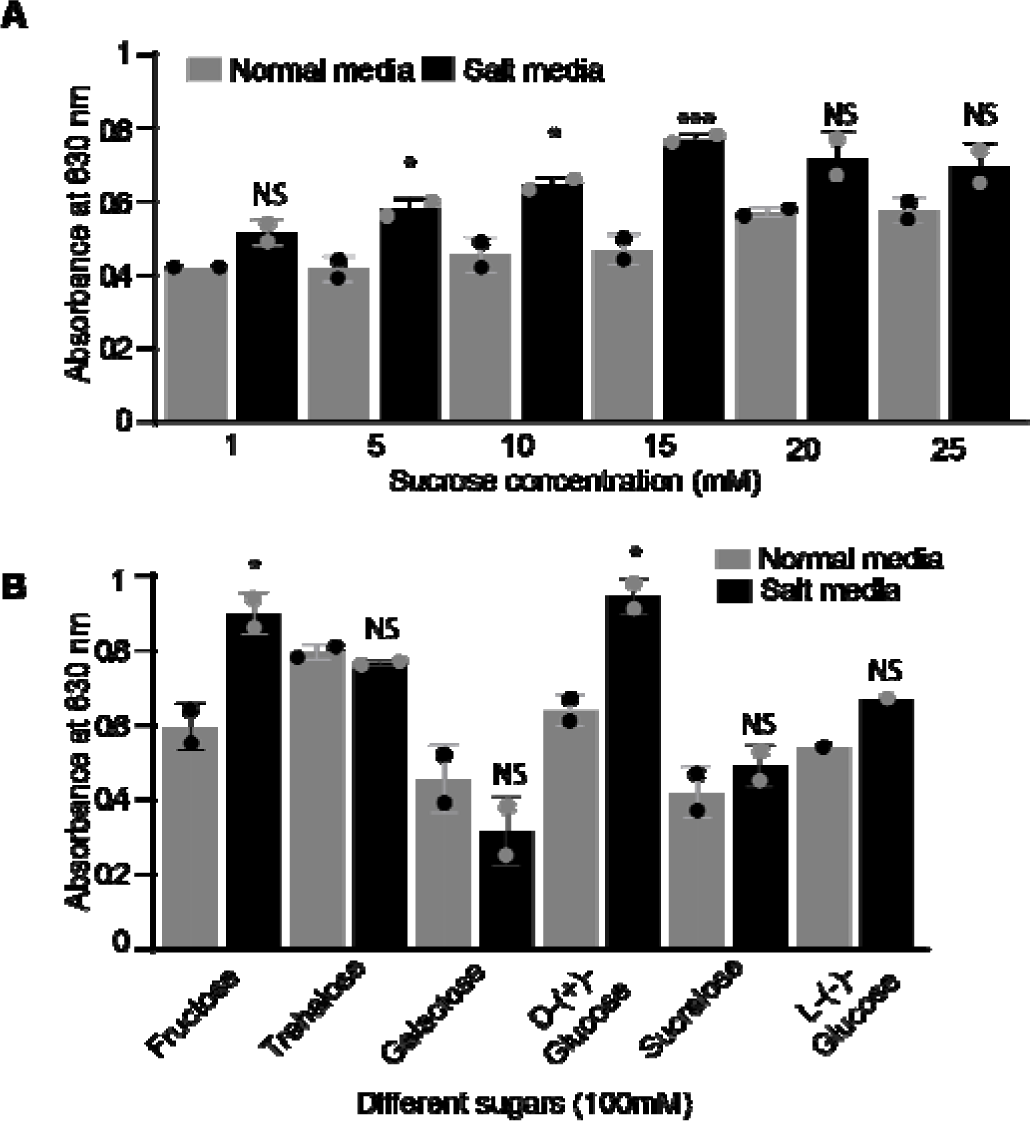
High NaCl fed flies show preference for selective sugars. (**A**) Absorbance values for different concentrations of sucrose (1, 5,10,15, 20 and 25mM) between high NaCl fed and standard media fed wildtype flies (*CsBz*). (**B**) Absorbance values for different sugars (fructose, trehalose, galactose, D-glucose, sucralose and L-glucose) at 100mM concentration between high NaCl fed and normal media fed flies. For **A** and **B**, n= 2 sets each concentration, 60 flies each set. Statistical analysis was performed using ANOVA Tukey’s multiple comparison test for obtaining P values: *p < 0.05, **p < 0.005 and ***p < 0.0005. For all graphs, error bars=SEM and NS is not significant.

We also tested different sugars to check if high NaCl fed flies prefer any particular sugar. We observed high consumption for other nutritive metabolizable sugars including100mM D-fructose and D-glucose (**Figure 3B**). No significant difference was found in the case of D-trehalose. Similarly, we did not see any differences in consumption for galactose, non-nutritive sugars-sucralose, and L-(-) glucose when tested at 100mM concentrations (**Figure 3B**). Also, we did not observe significant changes in spectrophotometry analysis at lower 10mM concentrations of any tested sugar (**Supplementary Figure 4C**), nor in feeding responses between high NaCl- and standard media-fed flies for the bitter compound caffeine (10mM concentration) (**Supplementary Figure 4D**). Our data suggest alteration in consumption of specific sugars (100mM concentration) in high NaCl-fed flies.

We also tested high NaCl fed flies for many other salts. We did not observe significant changes in feeding behavior between high NaCl- and standard media-fed flies (**Supplementary Figure 4E**) for different categories of salts. We tested 25 and 100mM concentrations of Sodium hydrogen carbonate, Di-sodium hydrogen O-phosphate, and 10 and 100mM concentrations of Magnesium chloride, and Potassium di-hydrogen O-phosphate.

### Role of neuronal activity in the peripheral and sweet LSO neurons of the pharynx in modulating high salt intake behavior under starvation

We next investigated the effect of neuronal activity in the peripheral sweet taste neurons on high salt intake behavior. First, we tested the *Gr5a* positive sugar receptor neurons (**Figure 4A**). *Gr5a* receptor is expressed only in the peripheral sweet neurons of the labellum (not in the neurons of the internal taste organs of the pharynx) [27]. To probe the role of *Gr5a* gustatory receptor neurons (GRNs) in mediating high salt intake we used flies with *Gr5a*-silenced neurons by expressing an active form of tetanus toxin (*Gr5a-GAL4>UAS-TNT*) and pre-exposed them to high NaCl diet. The expression of tetanus toxin makes neurons lose their activity and neurotransmitter release [28]. In our spectrophotometry and feeding assays, high NaCl fed test flies (*Gr5a-GAL4>UAS-TNT*) showed increased NaCl consumption (high absorbance values) and preference in feeding assays at 200mM NaCl and low NaCl (50mM) concentration (**Figure 4B and Supplementary Figure 5A**, black bars) when compared to standard media fed flies (**Figure 4B and Supplementary Figure 5A** - grey bars). High salt-fed flies showed high absorbance value and feeding preferences more than double at 50mM concentration (**Figure 4B and Supplementary Figure 5A**). Since, high salt (200mM) is an aversive cue, we also tested bitter compound caffeine (10mM) in our feeding assay. We did not observe any change in caffeine feeding responses when standard media fed *Gr5aGAL4>UAS-TNT* flies were compared to high salt-fed conditions (**Supplementary Figure 5D**). Both parental controls (*Gr5aGAL4/+* and *UAS-TNT/+*; **Figures 4C, D; Supplementary Figures 5B, C**) behaved as wildtype flies (**Figures 2B-D**) and showed increased feeding preferences and absorbance value in case of 200mM (high salt fed flies) but not at 50mM.

**FIGURE 4.**
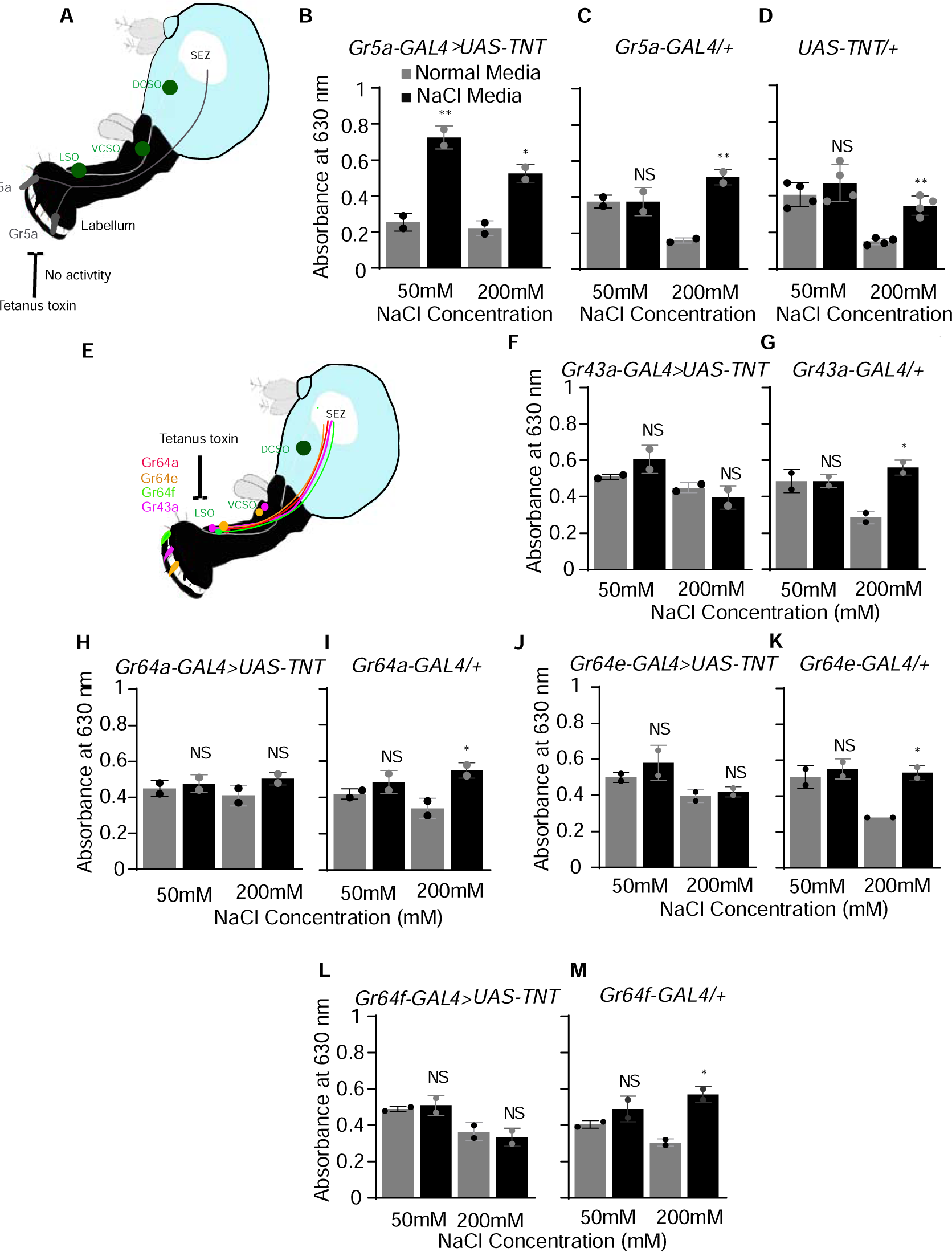
Role of neuronal activity in the peripheral and sweet LSO neurons of the pharynx in modulating high salt intake behavior. (**A**) Cartoon showing expression of *Gr5a*. (**B-D**) Comparison of mean absorbance values of flies after silencing neuronal activity of *Gr5a* sweet neurons through genetic expression of an active form of tetanus toxin (*Gr5a-GAL4>UAS-TNT*) with those of parental control flies (*Gr5a-GAL4/+ and UAS-TNT/+*). This analysis was carried out during a 2-h feeding assay in which flies consumed a mixture of NaCl and blue dye, with absorbance measured at 630nm. N= 2-4 sets each concentration, 60 flies each set. The graphs display absorbance values for low (50 mM) and high (200 mM) concentrations of NaCl after 24h of starvation. In all graphs, black bars represent responses of flies pre-exposed to high NaCl diet for 3 days compared with flies fed on normal fly media for 3 days (gray bars). Asterisks show significant differences between black vs grey for genotypes *Gr5a-GAL4>UAS-TNT*, *Gr5a-GAL4/+ and UAS-TNT/+*. (**E**) Cartoon showing expression of *Gr43a*, *Gr64a*, *Gr64e*, and *Gr64f* in LSO. (**F-M**) Mean absorbance values of flies after genetically manipulating neuronal activity of other sweet taste receptor neurons by expressing tetanus toxin (*UAS-TNT*) using *Gr43a-GAL4*, *Gr64a-GAL4*, *Gr64e-GAL4*, and *Gr64f-GAL4* drivers and their parental control flies (*Gr43a-GAL4/+*, *Gr64a-GAL4/+*, *Gr64e-GAL4/+*, and *Gr64f-GAL4/+*). N= 2 sets each concentration, 60 flies each set. Statistical analysis was performed using ANOVA Tukey’s multiple comparison test for obtaining P values: *p < 0.05, **p < 0.005 and ***p < 0.0005. For all graphs, error bars=SEM and NS is not significant.

Next, we looked at the consumption and feeding behavior of flies expressing other sweet taste receptors, including *Gr43a* (**Figure 4F and Supplementary Figure 5E**), *Gr64a*, *Gr64e,* and *Gr64f* (**Figures 4H, J, L and Supplementary Figures 5G, I, K**) after knocking their neuronal activity down by expressing tetanus toxin. Unlike *Gr5a,* which is only expressed in labellum neurons, these sweet taste receptors are expressed in LSO pharyngeal neurons (**Figure 4E) and have** an additional expression in the taste neurons of the labellum or in other regions of pharynx [29-32]. It has been shown that *Gr43a* and *Gr64e* co-express with other members of the sweet clade in LSO [33]. In case of silencing the neuronal activity of *Gr43a, Gr64a*, *Gr64e,* and *Gr64f* sweet LSO receptor neurons we did not observe change in the consumption or feeding preferences when 50 and 200mM NaCl concentrations were tested with standard media- and high NaCl media-fed flies (**Figures 4F, H, J, L and Supplementary Figures 5E, G, I, K**). Knock down of neuronal activity in these sweet LSO neurons lowered the feeding preferences back to normal, as seen in case of standard media-fed flies. The parental control flies (*UAS-TNT/+*, *Gr43aGAL4/+, Gr64aGAL4/+, Gr64eGAL4/+, Gr64fGAL4*/+ shown in **Figures 4D, G, I, K, M and Supplementary Figures 5C, F, H, J, L**) responded as wildtype flies (**Figures 2B-D**). These flies showed increased consumption and feeding preferences at 200mM concentration in case of high salt-fed flies but not at 50mM (between standard- and high NaCl-fed flies; grey and black bars). Our results suggest that functional and active LSO pharyngeal neurons under starvation (as seen in the case of *Gr5a>TNT* or wildtype flies; **Figures 2B-D, 4B and Supplementary Figure 5A**) regulate high NaCl intake.

### Role of neuronal activity in the peripheral and bitter LSO neurons of the pharynx in modulating high salt intake behavior under starvation

Our results suggest that high NaCl exposed *Gr5a*-silenced flies (only silenced peripheral labellum sugar-sensing neurons), consumed significantly more NaCl at both low (50mM) and high (200mM) concentrations compared to *Gr5a*-silenced flies fed on normal food. Notably, this phenotype was not observed in flies where pharyngeal sugar sensing neurons were silenced, suggesting neurons in the pharynx may play a role mediating the shift in preference observed following pre-exposure to high NaCl diet. We again questioned the mechanism by which this occurs, and tested flies by silencing bitter-neurons to see if bitter taste pathways are recruited in high salt aversion. It has already been shown [2] that high salt recruits two primary aversive taste pathways in mice by activating sour and bitter taste-sensing cells. Genetic silencing of sour and bitter pathways eliminate behavioral aversion to concentrated salt without impairing salt attraction. Mice lacking salt-aversion pathways exhibit unrestricted, continuous attraction even to tremendously high concentrations of NaCl. To test if the same is true in insects, we looked at the role of *Gr33a* positive bitter receptor neurons already known to detect the bitter compound caffeine. Since *Gr33a* does not express in LSO pharyngeal neurons [30, 34], we determined if silencing the activity of bitter neurons at the periphery (**Figure 5A**) would result in high NaCl feeding as seen in the case of *Gr5a>TNT* flies. We silenced the activity of bitter neurons (*Gr33a-GAL4>UAS-TNT*) by expressing tetanus toxin (**Figure 6B and Supplementary Figure 5A**) and found increased consumption and feeding preference in high NaCl-fed flies compared to standard media flies at 200mM concentration of NaCl (**Figure 5B and Supplementary Figure 6A**). The parental control flies (*Gr33aGAL4/+* and *UAS-TNT/+;* **Figures 5C, D and Supplementary Figures 6B, C)** showed similar absorbance values and feeding responses as wildtype flies (**Figures 2B-D**). We observed an increase in absorbance and feeding preferences only at 200mM in high salt fed flies. We did not find that *Gr33a*-silenced flies without high salt pre-exposure (standard media) consume large amounts of high salt (200mM) suggesting bitter pathways other than *Gr33a* are involved in aversive concentrations of NaCl (or high salt concentrations).

**FIGURE 5.**
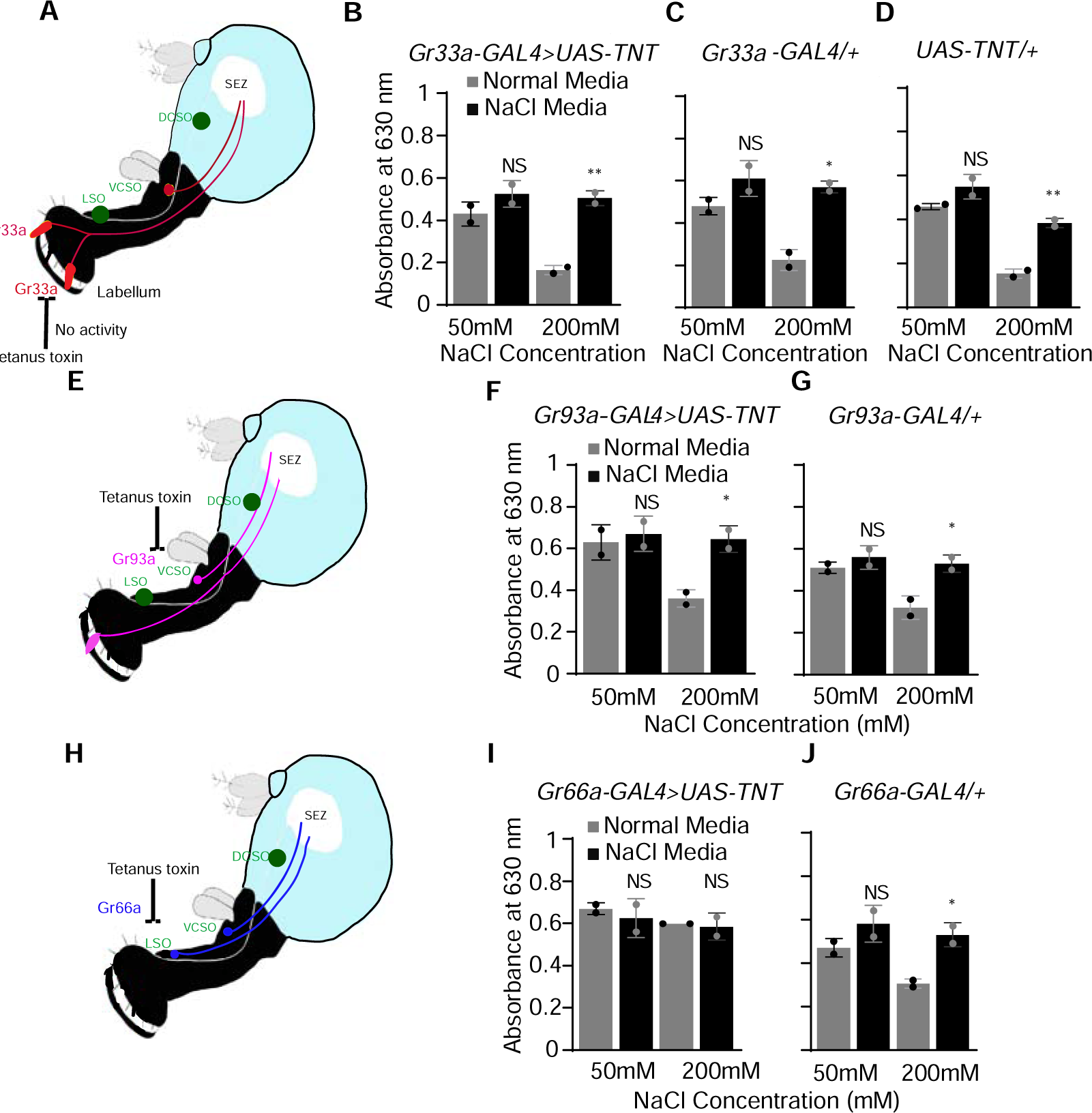
Role of neuronal activity in the peripheral and bitter LSO neurons of the pharynx in modulating high salt intake behavior. (**A**) Cartoon showing expression of *Gr33a* (**B-D**) Mean absorbance (spectrophotometry analysis) values of flies after silencing neuronal activity of bitter neurons by expressing an active form of tetanus toxin along with parental control flies (*Gr33a-GAL4>UAS-TNT, Gr33a-GAL4/+* and *UAS-TNT/+*) at the indicated NaCl concentrations after 24h of starvation. Asterisks show significant differences between *Gr33a-GAL4>UAS-TNT* flies fed on normal (grey bars) and high salt media (black bars). (**E)** Cartoon showing expression of *Gr93a*. (**F** and **G**) Spectrophotometry analysis of flies after silencing activity of *Gr93a* positive bitter neurons by expressing *UAS-TNT* (*Gr93a-GAL4>UAS-TNT*) (**F**) *along* with parental control flies (*Gr93a-GAL4/+*) (**G**). (**H**) Cartoon showing *Gr66a* expression. (**I** and **J**) Spectrophotometry analysis of flies after TNT-mediated silencing of *Gr66a* bitter neurons (*Gr66a-GAL4>UAS-TNT*) along with parental control (*Gr66a-GAL4/+)* (**I and J**) comparing normal media and high NaCl media feeding (gray vs black bars). N= 2 sets each concentration, 60 flies each set. Statistical analysis was performed using ANOVA Tukey’s multiple comparison test for obtaining P values: *p < 0.05, **p < 0.005 and ***p < 0.0005. For all graphs, error bars=SEM and NS is not significant.

**FIGURE 6.**
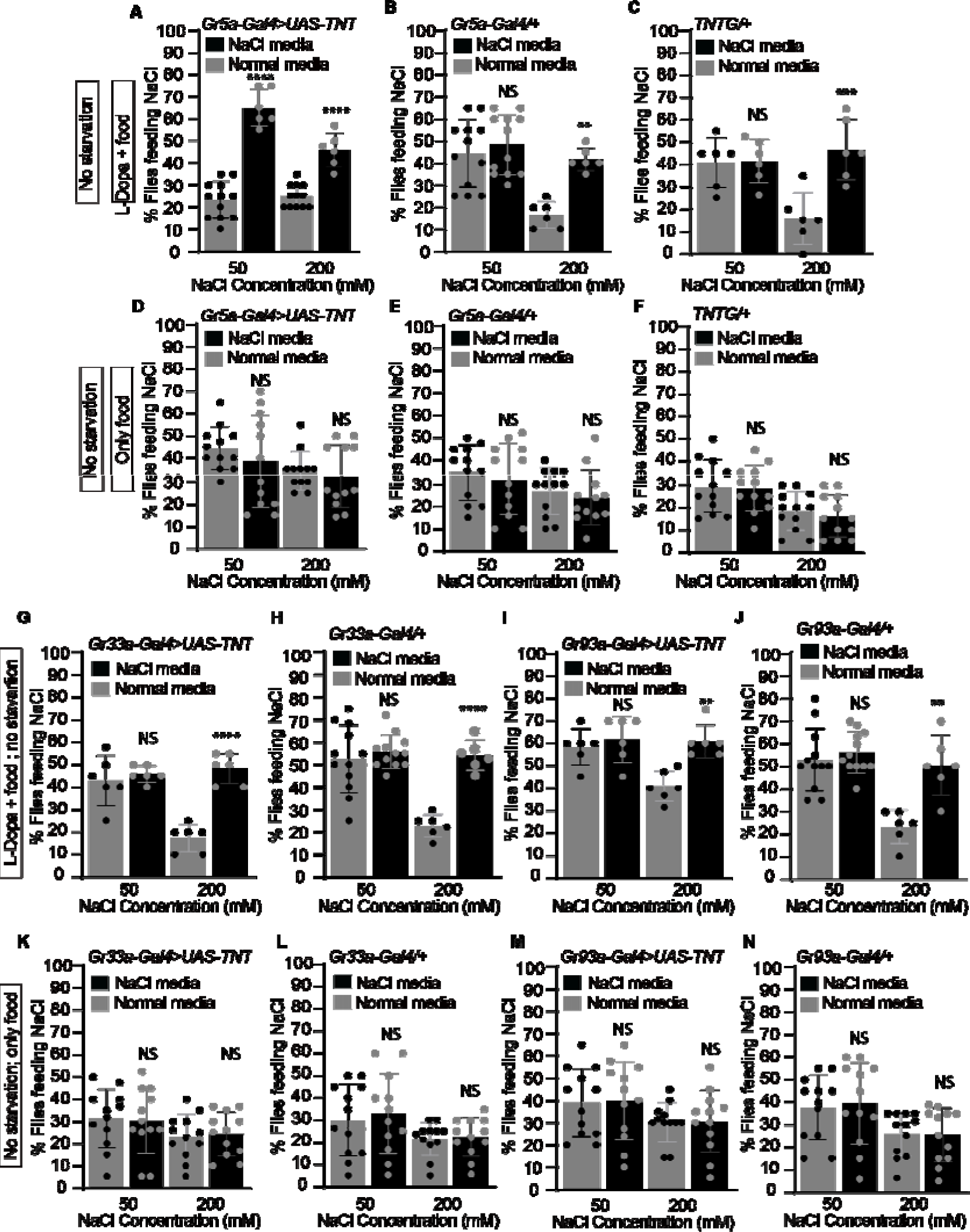
Role of Dopamine in modulating salt taste behavior. (**A-C**). Comparison of feeding preferences of flies with silenced neuronal activity of *Gr5a* sweet neurons through tetanus toxin expression (*Gr5a-GAL4>UAS-TNT*) and parental control flies (*Gr5a-GAL4/+ and UAS-TNT/+*) after L-Dopa feeding. In all graphs, black bars represent preferences of high NaCl fed flies compared with flies fed on normal fly media (grey bars). Asterisks show significant differences between black vs grey for genotypes *Gr5a-GAL4>UAS-TNT*, *Gr5a-GAL4/+ and UAS-TNT/+*. (**D-F**) Comparison of feeding preference of flies with silenced neuronal activity of *Gr5a* sweet neurons and parental control flies (*Gr5a-GAL4/+ and UAS-TNT/+*) after feeding on normal food (no L-dopa treatment). (**G-N**) Mean feeding preference of flies after genetically manipulating the neuronal activity of bitter taste receptor neurons with the tetanus toxin (*UAS-TNT*) using *Gr33a-GAL4* and *Gr93a-GAL4* and their parental control flies (*Gr33a-GAL4/+* and *Gr93a-GAL4/+*) with (**G-J**) and without L-Dopa treatment (**K-N**). For each bar, n=6-12 trails of 20 flies each (10 males and 10 females). Statistical analysis was performed using ANOVA Tukey’s multiple comparison test for obtaining P values: *p < 0.05, **p < 0.005 and ***p < 0.0005. For all graphs, error bars=SEM and NS is not significant.

Next, we looked at the feeding behavior of flies by selecting other classes of bitter receptor neurons, including *Gr93a,* that does not exist in LSO [20, 30, 35] (**Figures 5E, F**), and *Gr66a,* which is present in LSO [30, 36] (**Figures 5H, I and Supplementary Figures 6D, F**). We used tetanus toxin to knock down their neuronal activity (*Gr93a-GAL4>UAS-TNT* and *Gr66a-GAL4>UAS-TNT*). Since *Gr66a* is expressed in LSO pharyngeal neurons, silencing the neuronal activity of *Gr66a* receptor LSO neurons did not affect feeding preferences between standard- and high NaCl-fed media flies (**Figure 5I and Supplementary Figure 6F**). In case of the *Gr93a* (*Gr93a-GAL4>UAS-TNT*), we observed enhanced consumption and feeding preference (**Figure 5F and Supplementary Figure 6D**) at 200mM concentration similar to *Gr33a,* emphasizing again the role of active LSO neurons in regulating high salt intake in high NaCl-fed flies. The parental control flies (*Gr93aGAL4/+, Gr66aGAL4/+,* and *UAS-TNT**/+;*** **Figures 5D, G, J and Supplementary Figures 6C, E, G**) showed absorbance values and feeding responses as wildtype flies (**Figures 2B-D**).

The feeding preference and consumption at 200mM were found higher for *Gr66a>TNT* flies in case of standard media fed flies. We found weakened aversion at 200mM (compare standard media-fed flies grey bars only, **Figure 5I and Supplementary Figure 6F**), suggesting a possible role of *Gr66a* positive LSO bitter neurons in regulating high salt intake even under standard media condition.

### Genetic suppression of LSO pharyngeal neurons in high NaCl fed flies inhibits excessive salt intake

To again confirm and probe the role of LSO pharyngeal neurons in regulating high salt intake, next, we silenced pharyngeal LSO neurons specifically. For this, we used *Gr2a-GAL4* and *Poxn-GAL4* [30] as both show expression in the LSO region only (**Supplementary Figures 7A, B**, white arrow in right-hand side images). Silencing pharyngeal LSO neurons (*Gr2a-GAL4>UAS-TNT* and *Poxn-GAL4>UAS-TNT*) showed no significant changes in feeding behavior between high salt and standard media fed flies (**Supplementary Figures 7C, E,** left hand-side graphs). No significant differences were observed for low and high salt concentrations of NaCl even, in the case of spectrophotometry analysis (**Supplementary Figures 7D, F**, left hand-side graphs) at both concentrations. The parental control flies (*Gr2a-GAL4/+, poxn-GAL4/+* and *UAS-TNT/+;* **Supplementary Figures 7C-F**, right-hand side graphs; **Figure 4D and Supplementary Figure 5C**) showed absorbance values and feeding preferences as wildtype flies (**Figures 2B-D**).

Next, we tested *Drosophila Ir76b* receptor neurons. *Ir76b* is required for both high and low salt taste [13, 17, 37], and we confirmed its distribution in many LSO neurons (**Supplementary Figure 8a**, white arrow). We found no difference in feeding preferences between high salt and standard media fed flies (*Ir76b GAL4>UAS-TNT*; **Supplementary Figure 8B**) at 50mM and 200mM NaCl concentrations after silencing the neuronal activity of *Ir76b* neurons. *Ir76b GAL4>UAS-TNT* flies showed lower feeding responses at 50 and 200mM concentrations than parental flies (compare grey bars in **Supplementary Figures 8B, C**, **5C**). *Ir76b GAL4/+* flies behaved liked wild-type flies and showed increased feeding responses at 200mM when high salt-fed flies were compared to standard media-fed flies. Our behavioral results suggest that active LSO pharyngeal neurons are required and necessary to regulate high salt intake in flies pre-exposed to high salt under starvation condition. In the absence of activity in LSO neurons, flies show no difference in feeding behavior between standard media and high NaCl-fed conditions.

### Role of dopamine and activity in regulating high salt consumption

Recent studies have shown that starvation and L-dopa increase behavioral sensitivity to sucrose in flies [13, 17, 37-39]. To measure whether the activity of LSO pharyngeal neurons in high NaCl-fed flies gets modulated via dopamine signaling, we administered 3-(3,4-dihydroxyphenyl-2,5,6-d3)-L-alanine (L-Dopa) via standard food and measured feeding preferences. In this case, we did not starve flies in any of our experiments. Consistent with the results of starved flies, we found that high NaCl-fed flies (*Gr5a-GAL4>UAS-TNT*) showed increased feeding preferences at 200mM and 50mM NaCl concentrations (**Figure 6A**, black bars) after L-Dopa feeding (L-Dopa + food). Enhanced feeding was observed at 50mM NaCl concentration (**Figure 6A**, black bars) when compared with parental controls (*Gr5aGAL4/+* and *UAS-TNT/+*; **Figures 6B, C**) on high salt media (black bars). Under normal media fed conditions (no feeding on L-Dopa), we didn’t see any significant differences between standard and high salt-fed flies (*Gr5a-GAL4>UAS-TNT, Gr5aGAL4/+* and *UAS-TNT/+*) at 50 and 200mM (No starvation; **Figures 6D-F**).

Similarly, in case of *Gr33a-GAL4>UAS-TNT* (**Figure 6G**) and *Gr93a-GAL4>UAS-TNT* (**Figure 6I**) flies, we found high feeding preferences in high NaCl fed flies compared to standard media flies at 200mM (not at 50mM) concentration after feeding on L-Dopa. The parental control flies (*Gr33aGAL4/+, Gr93aGAL4/+* and *UAS-TNT**/+;*** **Figures 6C, H, J**) showed feeding preferences as wild-type flies (**Figure 2B**), where we observed increase in feeding preferences only at 200mM NaCl concentration (not at 50mM) between standard- and high salt-fed flies (grey and black bars). Under standard media fed condition (without L-Dopa), we did not see any significant differences between standard media- and high salt-fed flies (*Gr33a-GAL4>UAS-TNT, Gr93a-GAL4>UAS-TNT, Gr33aGAL4/+*, *Gr93aGAL4/+* and *UAS-TNT/+*) at 50 and 200mM (**Figures 6F, K-N**). Our results suggest modulation of salt taste behavior by both starvation or via dopamine in the presence of neuronal activity in LSO neurons.

## Discussion

Apart from peripheral taste cells, distinct internal taste organs are present in the pharynx of adult flies, namely LSO, VCSO (ventral cibarial sense organ), and DCSO (dorsal cibarial sense organ). After the initiation of food intake, the pharynx controls the ingestion of food and encourages only intake of appetitive food in flies. A recent receptor–to-neuron map of pharyngeal taste organs describes distinctive functional groupings of pharyngeal neurons [30]. While recent years have seen increased progress on peripheral salt coding in flies, the role of pharyngeal neurons in modulating salt intake remains unclear. Only recently, an inhibitory mechanism for suppressing high salt intake [24] and a role of single pair of pharyngeal neurons to reject high salt has been documented [25].

Calcium imaging experiments in the past suggested that *Gr5a-Gal4* sweet neurons mediate low salt attraction in insects [6] and that this driver labels additional non-sweet GRNs outside the sweet class [31]. Other groups proposed the role of Ionotropic receptor *Ir76b* [13, 37]. and *Gr66a* GRNs [6, 40] in mediating salt responses. *Gr64f* and *Ir94e* mediate attraction towards low salt and *Gr66a* and *Ppk23^glut^* drive avoidance to high salt concentrations [17], suggesting another layer of complexity of salt coding in flies. Recently, two groups showed the role of different Ir’s in mediating appetitive behavioral responses to salt [20] and normal avoidance of high monovalent salt concentrations [21]. In all these studies, the role of internal organs, including pharyngeal areas, was not tested for salt taste behavior.

Our results define the role of LSO pharyngeal neurons in regulating high salt intake and modulation of salt taste behavior. Our study suggests that increased dietary salt modulate and reshape salt curves to promote overconsumption of food in flies through LSO pharyngeal neurons in an activity- and state-dependent manner. We found that dopamine signaling also plays a role in this modulation. Multiple taste receptor neurons and pathways are independently involved in LSO neurons contributing to one output. When one of them is inhibited, we observed a partial reduction in increased salt responses. Our data suggest that genetic suppression of LSO neurons inhibits excessive salt intake, demonstrating role of this neuronal population in regulation of salt feeding behavior. Our data also suggest alteration in consumption of metabolizable sugars in high NaCl-fed flies, but not of non-metabolic sugars. Compare to fructose, which is detected by *Gr43a* [41], sucrose is detected by multiple receptors. Sucrose receptor abundance may help flies detect even the lower concentration of sucrose (less than 100mM; **Figure 3**).

Reducing activity specifically in LSO neurons, as seen with *Gr2a-GAL4* and *poxn-GAL4,* strengthens our observations that LSO neurons are necessary (**Supplementary Figure 7**). However, further experiments are required to prove if they are sufficient as well, as we could not assess whether activity in the LSO neurons alone would trigger the loss of salt aversion. As suggested in a recent study that salt taste is encoded by the combined activity of most of all GRN classes at the periphery [17], our results also hint that under starvation neurons of LSO region control and regulate high salt ingestion via many channels instead of one dedicated line. However, we do not know whether pre-exposure to high salt modulates higher brain areas, as the identity of high salt structures in the brain is still elusive (**Figure 8**).

Fruit flies, moths, and locusts abate food avoidance of certain bitter foods after prolonged exposure [42]. Dietary exposure to the unappealing but safe additives like camphor causes decline in camphor rejection. The long-term feeding on the camphor diet has been shown to result in reversible down regulation of TRPL (transient receptor potential-like) levels in the proboscis [37]. The state-dependent modulation of salt taste behavior mediated by *Ppk23^glut^* was shown under salt fed (3-day feeding with food containing 10mM NaCl) and salt deprivation condition [17]. Salt depletion in humans causes an increase in salt palatability [43]. The hunger signal overcomes aversive behavior to unappetizing foods [14]. Our study indicates a state-dependent role of internal pharyngeal taste neurons in the modulation of salt taste behavior (high and unappetizing concentration), which is least explored. We used a paradigm where 200mM NaCl (high and aversive concentration) was mixed with the standard food to pre-expose the flies to high salt condition for three successive days which is a different concentration than what Jaeger et al. used [17]. They utilized a concentration that is generally present in yeast (5-15%) added to standard fly media. This concentration of salt found in the fly food used to rear flies does not impact the PER or other feeding behavior responses as seen in our standard media condition results.

Mating modifies the feeding behavior in female *Drosophila* [44] and induces a salt appetite by increasing the gustatory response to sodium [26]. We also found that mated females show a higher feeding preference for salt (**Figure 1D**) suggesting, mated females need more sodium to invest in the progeny. Interestingly, dNPF and sNPF neuropeptides (short neuropeptide F) modulate multiple feeding-related behaviors, including control of food intake during starvation in flies [45-49]. During energy deficit conditions, animals become less selective in their food choices by enhancing their sensitivity to nutritious resources, such as sugar [18, 39, 50-53] suggesting the involvement of neuromodulatory cascades as critical mediators of state-dependent control [54-57]. Although we found dopamine-dependent modulation of salt taste behavior (**Figure 7**), the role of dNPF/sNPF signaling via LSO neurons in such a modulation calls for future investigation (**Figure 8**). Our data also suggest that high salt concentration (pre-feeding on high salt media) activates distinct taste pathways modulated by the hunger state as suggested by Devineni et al. [58] and drives opposing behaviors.

**FIGURE 7.**
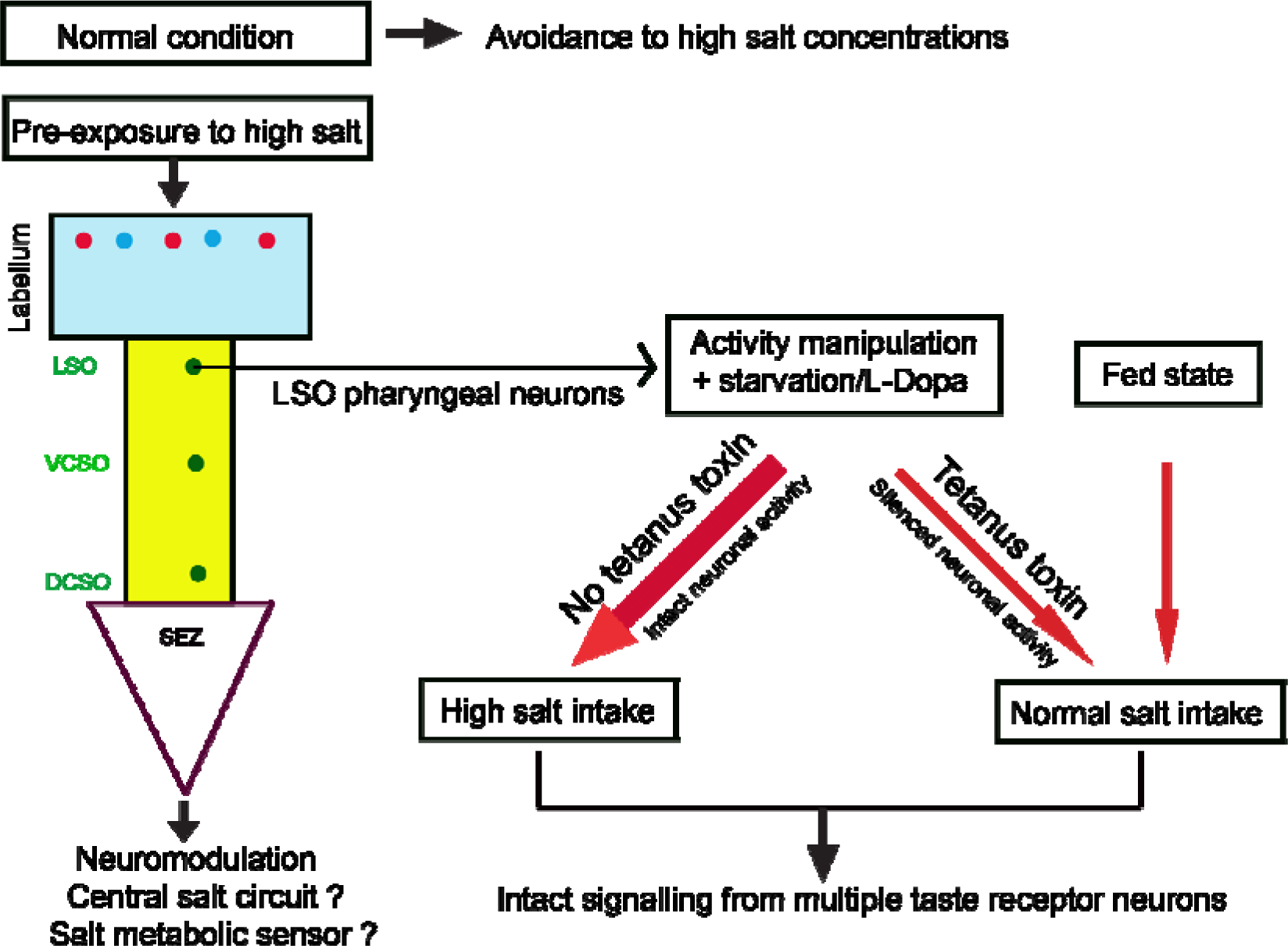
Working model of sodium intake. Three distinct internal taste organs are present in adult fly pharynx: the labral sense organ (LSO), ventral cibarial sense organ (VCSO), and dorsal cibarial sense organ (DCSO). Our study suggests that in “normal” conditions, flies have avoided high salt concentrations. If they were exposed to high salt AND either were starved afterwards, OR fed with L-Dopa, this avoidance is lost. The loss of avoidance requires intact signaling from several LSO neurons. Flies with intact functional LSO neurons under starvation or via dopamine signaling show increased feeding preference (thick red arrow) for low and high NaCl. The observed high NaCl feeding preferences get suppressed to normal feeding preferences in flies with reduced neuronal activity in LSO region as well as in fed state (red arrow) irrespective of what receptor they express (*Gr43a*, *Gr64a*, *Gr64e*, *Gr64f*, *Gr66a*, *Gr2a, Ir76b* and *poxn* all express in LSO) suggesting multiple pathways are involved in regulating high salt feeding in the hungry flies pre-fed on high salt. The identity of salt metabolic sensor in the brain and the role of central salt neurons causing high salt taste modulation is an open area of exploration.

The role of activity in modulating any taste has not been studied. It has been shown that acute alteration of the ventromedial hypothalamus (VMH) steroidogenic factor 1 (SF1) neurons in mice alters food intake [59] via changes in appetite and feeding-related behaviors. The study also found that SF1 neuron activity is sensitive to energy status. In our study, we observed changes in valence for NaCl taste in the presence of LSO neuronal activity under starvation conditions or via dopamine in high salt-fed flies. Our study defines an attractive population of neurons for regulating an essential aspect of feeding behavior in decision making and acute salt intake. How reduced neuronal activity in LSO neurons is connected to central circuits that control various aspects of feeding behavior under various internal states is yet to be determined. Our data suggest interesting roles of LSO pharyngeal neurons (sweet, bitter, and salt) where neuronal activity, and internal state play an important function in regulating salt intake. Such an analysis was missed in a study done by Chen et al. [23], where functional and behavioral studies on pharyngeal neurons in *poxn* mutants suggested avoidance of many aversive compounds, including high salt [23] by these neurons. Further identification and characterization of post synaptic high salt neuronal circuitry controlling ingestive behavior will significantly enhance our understanding of how internal state modifies perception and behavior towards unappetizing salt concentrations (**Figure 8**).

Our results suggest that pharyngeal taste organs like LSO play an essential role in sodium homeostasis. Neurons like LSO in the pharynx present a regulatory system that has evolved to fine-tune calorie intake with energy metabolism. Interestingly, consuming sodium can drive the desire to eat even more in many species, including humans. Examination of molecular and behavioral impact of appetitive cues like sugar or low salt/high salt on the brain’s reward circuitry [60] leads to overeating and metabolic issues are required to study the underlying mechanisms that drive changes in the neural activity. Answering this question may open up avenues to help people with health issues related to high salt consumption to eat less sodium in their diets.

## Materials and Methods

### Fly stocks

*CsBz* and *w^1118^* (from NCBS, Bangalore), *UAS-TNT* (BL 28838), *Gr5a-GAL4* (BL 57592), *Gr33a-GAL4* (BL 57623), *Ir76b-GAL4* (BL 51311), *Gr43a-GAL4* (BL 57636), *Gr64f-GAL4* (BL 57669), *Gr64a-GAL4* (BL 57661), *Gr64e-GAL4* (BL 57667), *Gr66a-GAL4* (BL 57670), *Gr93a-GAL4* (BL 57679), *Gr2a-GAL4* (BL 57590), *UAS-mcd8GFP* (BL 5137) and *poxn*-GAL4 (Bl. 66685) were obtained from the *Drosophila* Bloomington Stock Center. *Drosophila* stocks were reared on standard cornmeal dextrose medium at 25^0^ C unless specified otherwise.

The *Drosophila* media composition used was (for 1 liter of media) - corn flour (80g), D-glucose (20g, SRL-Cat. no.50-99-7), sugar (40g), agar (8g, SRL-Cat. no. 9002-18-0), yeast powder (15g, SRL-REF-34266), propionic acid (4ml, SRL, Cat. no. 79-09-4), TEGO (1.25 g in 3ml of ethanol, Fisher Scientific, Cat. no. 99-76-3), and orthophosphoric acid (600ul, SRL, Cat. no.7664-38-2).

### Feeding behavior assays

For feeding assays, flies of the required genotype were raised from eggs to adults at 25^0^ C. Flies were sorted in vials of 10 males and 10 females (20 flies/vial) upon eclosion and maintained at 25^0^ C for 3 days on fresh media (standard fly food or high NaCl media). Following starvation of flies for 24 hrs at 25^0^ C, they were tested for feeding behavior as described previously [38, 61], and abdominal coloration was scored as positive if there was any pink or red eating (red or pink-feeding on salt or taste compound; white-no feeding; blue-feeding on agar and water; purple-feeding on both red and blue). Purple was not scored in any of the experiments. 60X15mm feeding plates from Tarsons (India) were used for the assay. In these assays, flies were provided with red dots with taste compounds and blue spots as control containing dye, water and agar.

% flies feeding on taste compound was calculated as follows: First, % flies feeding on red or blue **(Supplementary Figure 1A)** for each plate was calculated. The Mean of % flies feeding for 6 or more plates was taken as a final value.

### Tarsal Proboscis extension reflex (PER) assay

For PER experiments, flies were tested as described previously [38]. Flies were collected after 2-3 days of eclosion and kept on standard food for 2 days. Both male and female flies were used for the PER assay. The flies were starved for 24 hrs in vials with water-saturated (4ml) tissue papers. Prior to the PER experiment, flies were immobilized by cooling on ice for at least 15 minutes and then mounted using nail polish, vertical aspect up, on glass slides (76mm X 26mm X 1mm from Borosil). Flies were allowed to recover in a moist chamber (plastic box with wet tissues) for at least 2 hrs prior to testing. Tastant solutions prepared in water were applied to tarsi via a drop extruded using 2ul pipette. Before testing the taste solutions, flies were allowed to drink water *ad libitum*. Flies not responding to water were excluded before the assay. Flies satiated with water were then tested with NaCl or other tastant solutions. Ingestion of any tastant solutions was not permitted, and, following each tastant application, flies were retested with water as a negative control. Each fly was tested five times with each tastant solution stimulus. The interval between consecutive tastant solution applications was at least 2-3 min to minimize adaptation. Flies showing three or more proboscis extensions were considered responders.

More than 50 flies were tested in batches for all PER experiments, and the percentage of responders was calculated for each set. Graphs depict mean responses, and error bars indicate SEM (Standard error of the mean).

### Chemicals

Sugars used were all obtained from Sigma Aldrich-Sucrose (57-50-1), Fructose (57-48-7), Trehalose (6138-23-4), Sucralose (56038-13-2), Galactose (59-23-4), L-(-) Glucose (Sigma –921-60-8) and D-(+)-Glucose (50-99-7). The other compounds used in the study-NaCl salt (Fisher Scientific-7647-14-5) which was of 99.9% purity; Caffeine (Sigma-Aldrich # 58-08-2); Blue dye-Indigo carmine (Sigma: 860-22-0); Red dye-Sulforhodamine B (Sigma-3520-42-1); Potassium Di Hydrogen orthophosphate (Fisher Scientific -7778-77-0); Magnesium Chloride (Fisher Scientific-7791-18-6); Di Sodium Hydrogen O-phosphate (Fisher Scientific-7558-79-4); Sodium Hydrogen carbonate (Fisher Scientific-144-55-8); Potassium Chloride (Fisher Scientific-7447-40-7) and Calcium chloride (Fisher Scientific-10043-52-4).

### Spectrophotometry analysis

After eclosion, 3-days old flies were sorted (10 males and 10 females) into batches on standard media which served as a control (X 6 vials). Similarly, 3 days old flies were separated as 10 males and 10 females on different salt concentrations (10, 50, 100, 200 and 500mM NaCl) mixed with normal fly food (X 6 vials). Flies were kept on these media conditions for 3 days as we did for feeding assay in Figure 3B.

For spectrophotometry analysis: after feeding assays, total 120 flies were divided into two sets (2 sets X 60 flies each) for each concentration. Later, flies (all the flies were used in the assay in an unbiased manner) were put in 2ml Eppendorf tubes in 70% ethanol (60 flies in one Eppendorf). Flies were first crushed in 150ul of 70% ethanol and then 150ul 70% ethanol was added to crush them more. After crushing, 200 ul of double distilled water (to get the content sticking to pestle and the wall of the Eppendorf tube) was added in the same soup and centrifugation was done at 3000 rpm for 15 minutes. After centrifugation, pellet was discarded and supernatant was transferred in the fresh Eppendorf’s. To do the spectrophotometry analysis, supernatant was further diluted with double distilled water to make up the total final volume of 650ul in the cuvette. Spectrophotometry analysis was done at 630 nm wavelength. One reading was taken for each sample and only the mean values were plotted. Spectrophotometer used was Perkin Elmer, lambda 35 UV/VIS Spectrometer.

### Immunohistochemistry

Immunohistochemistry for labellum was performed as mentioned before [38]. After anesthetizing flies on ice, labellum was dissected in chilled 1X PBS and kept for fixing (30 min) in 4% paraformaldehyde (0.3% Triton X-100) at room temperature. After washes with PBST (PBS with 0.3% TritonX-100), samples were blocked with 5% normal goat serum in PBST. Samples were incubated in appropriate primary antibody solutions overnight at 4^0^C. Primary antibody used was rabbit anti-GFP (1:1000, Invitrogen, catalog no. A11122) and secondary antibody used was Alexa Fluor 488 goat anti-rabbit immunoglobulin G (IgG) (1:200, Invitrogen).

### L-Dopa feeding experiments

For experiments in Figure 7, L-Dopa (Sigma-Aldrich, CAS no. - 53587-29-4) was first dissolved in water (5mg/ml) [38]. Then freshly prepared solution was spread on fly food. After this, flies were maintained on this L-Dopa food for 3 days in the dark at 25^0^ C incubators. The medium was changed once after 24 hr and freshly made L-Dopa was added to the media. Control flies were fed on normal fly food without any L-Dopa. No starvation was involved in these experiments.

## Microscopy used for image analysis and video recording

### GFP Imaging

Adult labellum was mounted in 70% glycerol in PBST after immunohistochemistry. Samples were analyzed and GFP fluorescence was visualized using a confocal microscope (Leica TCS SP5 II), and image stacks were generated acquired at 0.5 micron optical sections. Olympus SZX10 dual tube microscope was used for generating videos and Olympus SZ61 stereomicroscopes for doing the general fly pushing. Images were processed using ImageJ, Adobe Photoshop, and Illustrator software.

### Statistical analyses

Unless otherwise stated, all the results from behavioral experiments were analyzed for statistical significance with parametric ANOVA followed by post hoc Tukey multiple comparisons test for obtaining the p-values. We used student’s t-test for Supplementary Figure 4D and 5D.

## Supporting information

Supplemental media file 1

Supplemental media file 2

## Acknowledgements

We are grateful to PK lab members for helpful comments on the manuscript. We thank National Centre for Biological Sciences, Bangalore and Bloomington Drosophila Stock Center, USA for the fly stocks.

## Authors contributions

Conceptualization, P.K.; Methodology, T.T. and P.K.; Investigation, S.K, S.S, S.K, R.K; Validation, S.K., R.K and P.K.; Formal Analysis, S.K., R.K., D.E.R.L. and P.K.; Writing – Original Draft, P.K.; Writing – Review & Editing, S.K, R.K., T.T., D.E.R.L. and P.K..; Visualization, S.K, R.K. and P.K.; Figures Preparation, S.K. and P.K.; Supervision, P.K.; Funding Acquisition, P.K.

## Ethics statement

*Drosophila* strains used in the study was reviewed and approved by the institutional ethical committee.

## Data availability statement

The datasets generated during and/or analyzed during the current study are available from the corresponding author on reasonable request.

## Supplementary Material

**SUPPLEMENTARY FIGURE 1**

**Dye switching experiment**. (**A**) Graph showing mean feeding preference of flies after switching the dyes. Grey bars represent feeding preferences of flies for different concentrations of NaCl after 24 h of starvation. Salt was added to blue dye instead of red dye in these experiments. N= 18 plates, 20 flies each plate (10 males and 10 females). (**B**) Mean of positive PER responses in flies tested with 100mM sucrose each time before applying NaCl and by the end of the experiment in tarsal PER assay. N= 62 flies. Statistical analysis was performed using ANOVA Tukey’s multiple comparison test for obtaining P values: *p < 0.05, **p < 0.005 and ***p < 0.0005. For all graphs, error bars=SEM and NS is not significant.

**SUPPLEMENTARY FIGURE 2**

**Mean feeding preference of wildtype flies (*CsBz*) tested with various concentrations of NaCl on two different media conditions**. (**A**) Flies pre-exposed to high salt diet (black bars) and normal media for 1 day (grey bars), (**B**) and 2 days followed by 24hrs starvation condition. For each bar, n=6 trails of 20 flies each (10 males and 10 females). Statistical analysis was performed using ANOVA Tukey’s multiple comparison test for obtaining P values: *p < 0.05, **p < 0.005 and ***p < 0.0005. For all graphs, error bars=SEM and NS is not significant.

**SUPPLEMENTARY FIGURE 3**

**Flies show no difference in feeding preferences under fed state**. (**A**) % mean feeding preferences of wildtype flies pre-exposed to high salt diet (black bars) and normal media for 3 days (grey bars) under no starvation condition. For each bar, n=6 trails of 20 flies each (10 males and 10 females). (**B**) KCl dose response curve of normal media (grey bars) and high salt media fed flies (black bars). N=6-12 plates, 20 flies each plate (10 males and 10 females). Statistical analysis was performed using ANOVA Tukey’s multiple comparison test for obtaining P values: *p < 0.05, **p < 0.005 and ***p < 0.0005. For all graphs, error bars=SEM and NS is not significant.

**SUPPLEMENTARY FIGURE 4**

**Flies show no change in feeding preferences for high sucrose concentration and other salts**. (**A and B**) Absorbance values and mean feeding preference of wildtype (*CsBz*) flies for high concentrations of sucrose (50,100 and 200mM). (**C**) Absorbance values for different sugars (fructose, trehalose, galactose, D-glucose, sucralose and L-glucose) at 10mM concentration between high NaCl and normal media fed flies. (**D**) Mean feeding preference of wildtype (*CsBz*) flies for caffeine (10mM). In all graphs, black bars represent responses of flies pre-exposed to high NaCl diet compared to grey bars (normal fly media fed flies). (**E**) Mean feeding preference of wildtype (*CsBz*) flies at the indicated concentrations of other salts - sodium hydrogen carbonate (25 and 100mM), Di-sodium hydrogen O-phosphate (25 and 100mM), Magnesium Chloride (10mM and 100mM and potassium di hydrogen O-phosphate (10mM and 100mM). For absorbance assays in **A** and **C**, n= 2 sets each concentration, 60 flies each set. For feeding assays in **B**, **D** and **E**, N=6 plates, 20 flies each plate (10 males and 10 females). Statistical analysis was performed using ANOVA Tukey’s multiple comparison test for obtaining P values: *p < 0.05, **p < 0.005 and ***p < 0.0005. For all graphs, error bars=SEM and NS is not significant.

**SUPPLEMENTARY FIGURE 5**

**Role of neuronal activity in the peripheral and sweet LSO neurons of the pharynx in modulating high salt intake behavior under starvation condition**. (**A**) Mean feeding preference of *Gr5a-GAL4>UAS-TNT* flies after silencing neuronal activity of peripheral *Gr5a* sweet neurons by expressing tetanus toxin at low (50mM) and high (200 mM) NaCL concentrations after 24h of starvation. In all graphs, black bars represent responses of high NaCl-fed flies compared with normal media-fed flies (grey bars). Asterisks show significant differences between *Gr5a-GAL4>UAS-TNT* flies (black vs grey bars). (**B** and **C**) Mean feeding preference of parental flies (*Gr5a-GAL4/+* and *UAS-TNT/+).* (**D**) Mean feeding preference of *Gr5a-GAL4>UAS-TNT* flies (both normal media and salt pre-exposed) for caffeine (10mM). (**E**, **G**, **I**, and **K**) Mean feeding preference of flies after genetically manipulating the neuronal activity of other sweet taste receptor neurons by expressing tetanus toxin (*UAS-TNT*) using *Gr43a-GAL4*, *Gr64a-GAL4*, *Gr64e-GAL4*, and *Gr64f-GAL4* drivers, respectively. (**F**, **H**, **J,** and **L**) Mean feeding preference of parental flies (*Gr43a-GAL4/+, Gr64a-GAL4/+, Gr64e-GAL4/+* and *Gr64f-GAL4/+)* for 50 and 200 mM NaCl. For each graph, each bar, n=6-12 trails of 20 flies for each concentration (10 males and 10 females). Student’s t-test in was used for D otherwise statistical analysis was performed using ANOVA Tukey’s multiple comparison test for obtaining P values: *p < 0.05, **p < 0.005 and ***p < 0.0005. For all graphs, error bars=SEM and NS is not significant.

**SUPPLEMENTARY FIGURE 6**

**Role of neuronal activity in the peripheral and bitter LSO neurons of the pharynx in modulating high salt intake behavior under starvation condition**. (**A**) Mean feeding preference of flies after genetically silencing activity of peripheral bitter neurons by expressing tetanus toxin (*Gr33a-GAL4>UAS-TNT*) at the indicated concentrations of NaCl after 24hrs of starvation. Asterisks show significant differences between *Gr33a-GAL4>UAS-TNT* flies fed on normal (grey bars) and salt media (black bars). (**B** and **C**) Mean feeding preference of parental control flies (*Gr33a-GAL4/+* and *UAS-TNT/+*). (**D** and **F**) Mean feeding preference of flies after silencing activity of other bitter neurons (*Gr93a* and *Gr66a*) by expressing *UAS-TNT* (**D**, *Gr93a-GAL4>UAS-TNT* and **F**, *Gr66a-GAL4>UAS-TNT*) between normal media and high NaCl media exposed flies (grey vs black bars). (**E** and **G**) Mean feeding preference of parental control flies (*Gr93a-GAL4/+* and *Gr66a-GAL4/+*). For each graph, each bar, n=6-12 trails of 20 flies for each concentration (10 males and 10 females). Student’s t-test in was used for D otherwise statistical analysis was performed using ANOVA Tukey’s multiple comparison test for obtaining P values: *p < 0.05, **p < 0.005 and ***p < 0.0005. For all graphs, error bars=SEM and NS is not significant.

**SUPPLEMENTARY FIGURE 7**

**Silencing LSO pharyngeal neurons shows no increase in feeding preference for NaCl in high salt-fed flies.** (**A**) Expression pattern of *Gr2a-GAL4* labeled by *UAS-mCD8GFP* in the LSO pharyngeal neurons (white arrow in right panel). (**B**) Expression pattern of *poxn-GAL4* marked by *UAS-mCD8GFP* in the LSO pharyngeal neurons (white arrow in right panel). (**C**) Mean feeding preference of flies after silencing neuronal activity in the *Gr*2a (*Gr2a-GAL4>UAS-TNT* and parental control *Gr2a-GAL4/+*) positive LSO neurons tested with 50mM and 200mM NaCl (between normal media- and high salt media-fed flies; compare grey vs black bars). (**D**) Spectrophotometry analysis of normal and high salt media fed *Gr2a-GAL4>UAS-TNT* and *Gr2a-GAL4/+* flies. (**E**) Feeding preference of flies after silencing neuronal activity in the *poxn* (*poxn-GAL4>UAS-TNT* and parental control *poxn-GAL4/+*) positive LSO neurons (compare grey vs black) tested with 50mM and 200mM NaCl. (**F**) Spectrophotometry analysis of normal media and high salt media *poxn-GAL4>UAS-TNT* and *poxn-GAL4/+* flies. For **C** and **E**, n=6-12 plates, 20 flies each plate. For spectrophotometry analysis (**D** and **F**), n= 2 sets each concentration, 60 flies each set. Statistical analysis was performed using ANOVA Tukey’s multiple comparison test for obtaining P values: *p < 0.05, **p < 0.005 and ***p < 0.0005. For all graphs, error bars=SEM and NS is not significant.

**SUPPLEMENTARY FIGURE 8**

**Silencing *Ir76b* LSO pharyngeal neurons showed no increased feeding preference for NaCl in high salt pre-exposed flies.** (**A**) Expression pattern of *Ir76b-GAL4* labeled with *UAS-mCD8GFP* in the LSO pharyngeal neurons (white arrow). (**B**) Mean feeding response of *Ir76b-GAL4>UAS-TNT* flies compared between normal media- and high salt media-fed flies. (**C**) Mean feeding preference of *Ir76b-GAL4/+* parental flies. Statistical analysis was performed using ANOVA Tukey’s multiple comparison test for obtaining P values: *p < 0.05, **p < 0.005 and ***p < 0.0005. For all graphs, error bars=SEM and NS is not significant.

**Supplementary movie 1 for Figure 1**.

Tarsal PER - Fly showing extension of proboscis when 50mM NaCl was presented

**Supplementary movie 2 for Figure 1**.

Tarsal PER - Fly showing no extension of proboscis when 200mM NaCl was presented.

## Abbreviations

ANOVA: analysis of variance
DCSO: dorsal cibarial sense organ
DEG; dNPF; *dpr* locus: defective proboscis extension response
ENaC: epithelial sodium channel
GFP; Gr; GRNs: gustatory receptor neurons
IgG: immunoglobulin G
Ir; KCl; L-Dopa: 3-(3,4-dihydroxyphenyl-2,5,6-d3)-L-alanine
LSO: labral sense organ
NaCl: Sodium Chloride
PER: Proboscis Extension Reflex
Poxn; ppk: *Pickpocket*
SEM: Standard error of the mean
SF1: steroidogenic factor 1
sNPF; TEGO; TMC-1: *trans*-membrane channels like
TNT; TRCs: Taste receptor cells
TRPL: transient receptor potential-like
VCSO: ventral cibarial sense organ
VMH: ventromedial hypothalamus.

