## Supplemental tables 1-8 for "Activity and state-dependent modulation of salt taste behavior via pharyngeal neurons in *Drosophila melanogaster*"

**Corresponding author**

Running title: Modulation of salt taste behavior in *Drosophila*

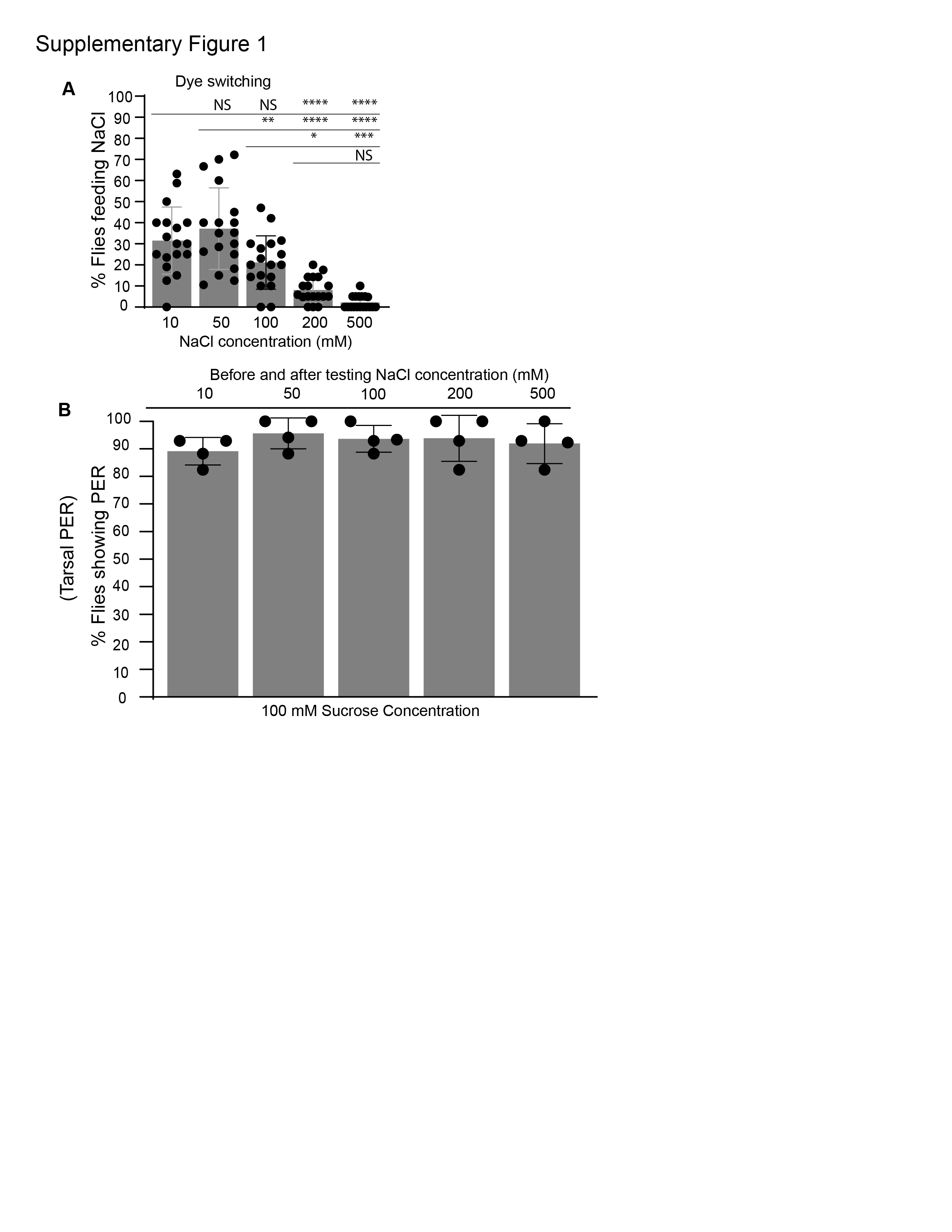

**SUPPLEMENTARY FIGURE 1**

**Dye switching experiment**. (**A**) Graph showing mean feeding preference of flies after switching the dyes. Grey bars represent feeding preferences of flies for different concentrations of NaCl after 24 h of starvation. Salt was added to blue dye instead of red dye in these experiments. N= 18 plates, 20 flies each plate (10 males and 10 females). (**B**) Mean of positive PER responses in flies tested with 100mM sucrose each time before applying NaCl and by the end of the experiment in tarsal PER assay. N= 62 flies. Statistical analysis was performed using ANOVA Tukey’s multiple comparison test for obtaining P values: *p < 0.05, **p < 0.005 and ***p < 0.0005. For all graphs, error bars=SEM and NS is not significant.

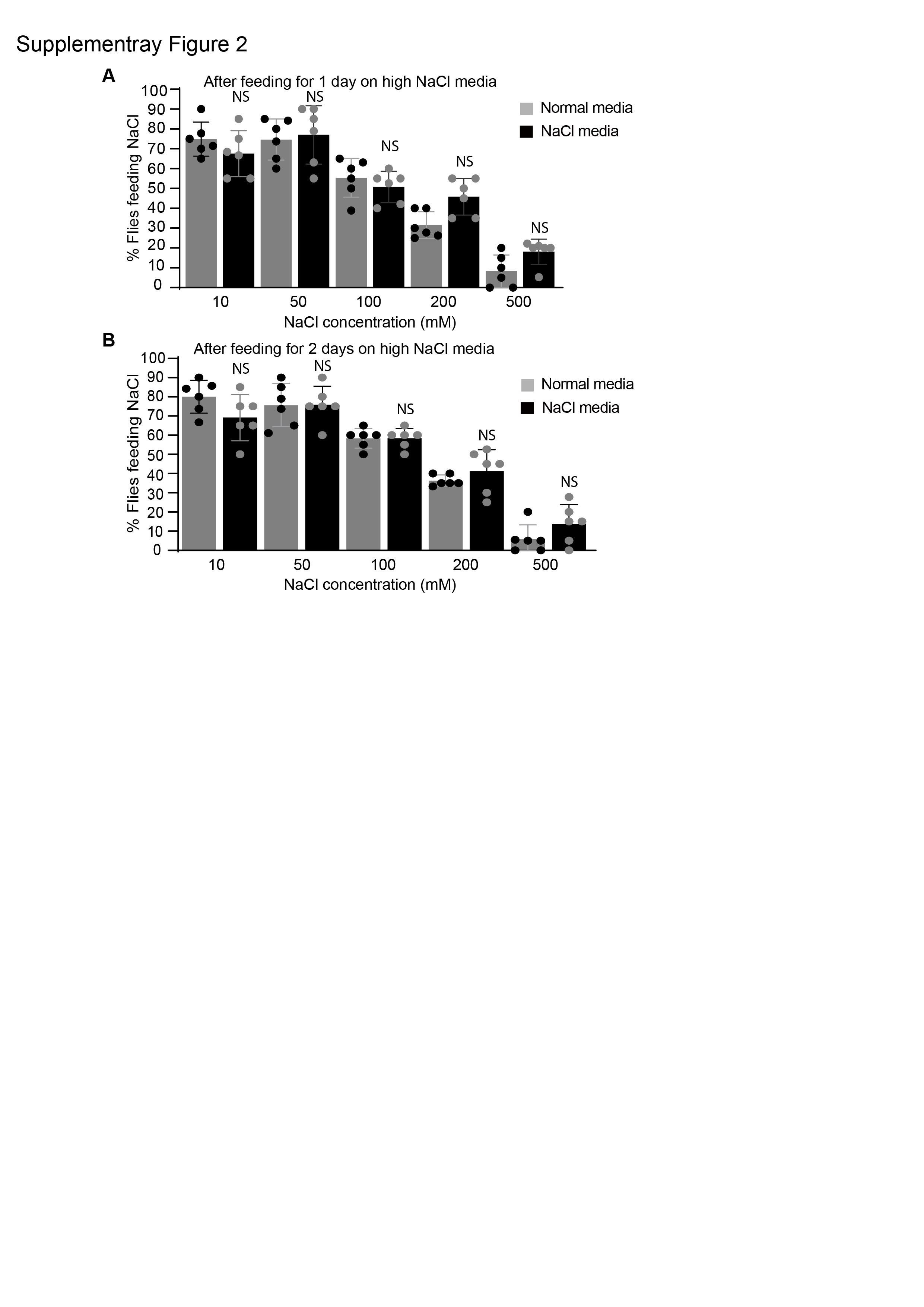

**SUPPLEMENTARY FIGURE 2**

**Mean feeding preference of wildtype flies (*CsBz*) tested with various concentrations of NaCl on two different media conditions**. (**A**) Flies pre-exposed to high salt diet (black bars) and normal media for 1 day (grey bars), (**B**) and 2 days followed by 24hrs starvation condition. For each bar, n=6 trails of 20 flies each (10 males and 10 females). Statistical analysis was performed using ANOVA Tukey’s multiple comparison test for obtaining P values: *p < 0.05, **p < 0.005 and ***p < 0.0005. For all graphs, error bars=SEM and NS is not significant.

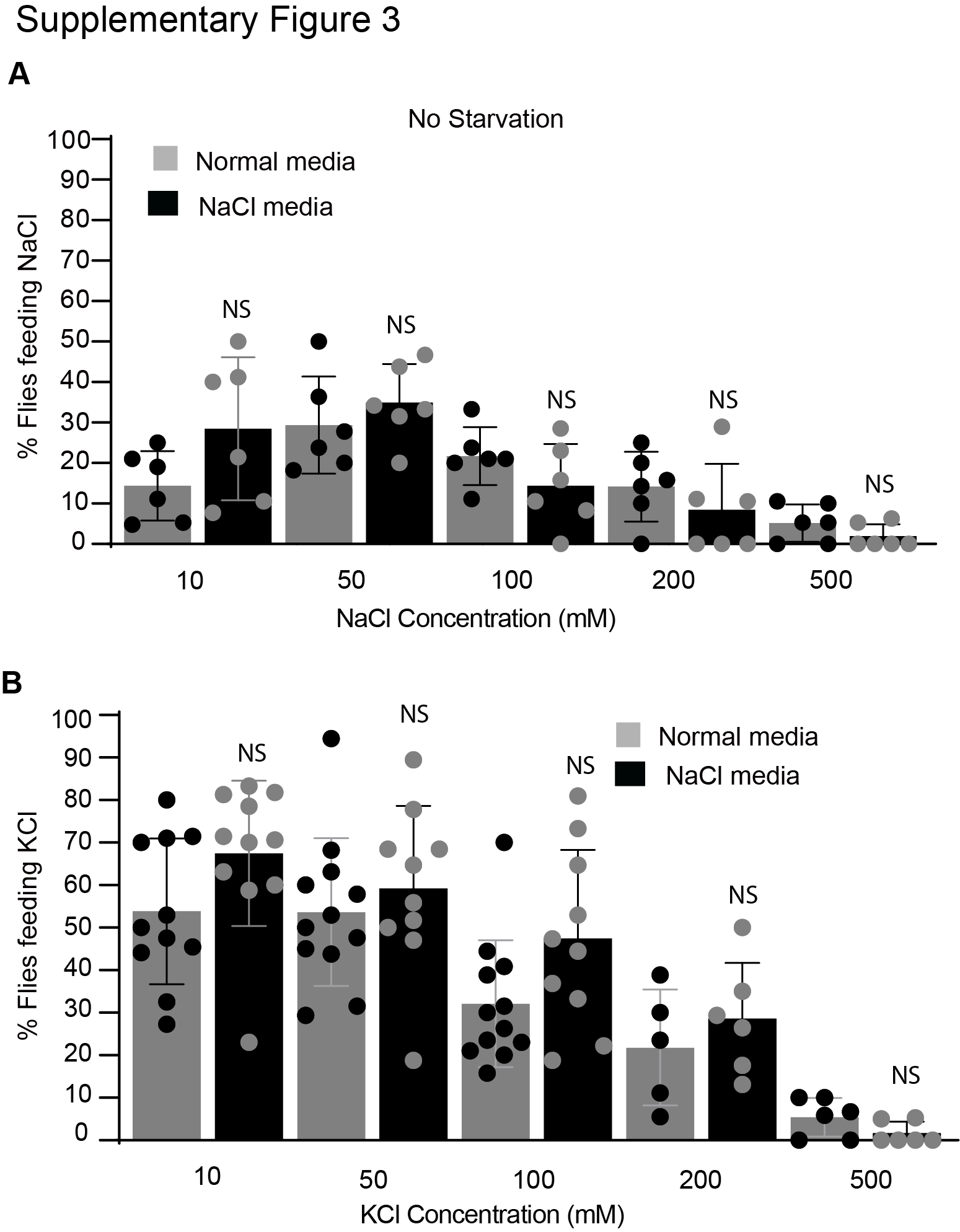

**SUPPLEMENTARY FIGURE 3**

**Flies show no difference in feeding preferences under fed state**. (**A**) % mean feeding preferences of wildtype flies pre-exposed to high salt diet (black bars) and normal media for 3 days (grey bars) under no starvation condition. For each bar, n=6 trails of 20 flies each (10 males and 10 females). (**B**) KCl dose response curve of normal media (grey bars) and high salt media fed flies (black bars). N=6-12 plates, 20 flies each plate (10 males and 10 females). Statistical analysis was performed using ANOVA Tukey’s multiple comparison test for obtaining P values: *p < 0.05, **p < 0.005 and ***p < 0.0005. For all graphs, error bars=SEM and NS is not significant.

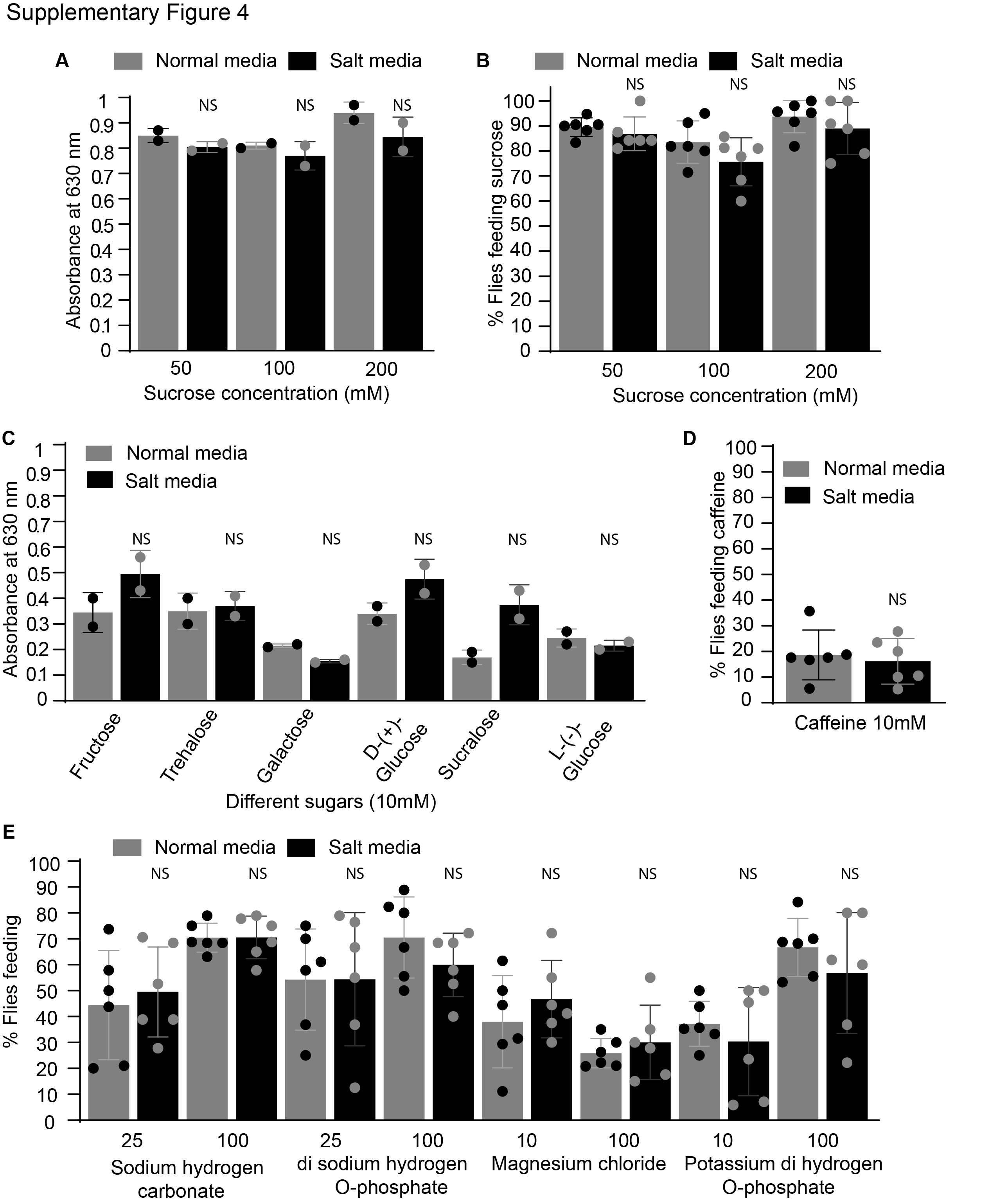

**SUPPLEMENTARY FIGURE 4**

**Flies show no change in feeding preferences for high sucrose concentration and other salts**. (**A and B**) Absorbance values and mean feeding preference of wildtype (*CsBz*) flies for high concentrations of sucrose (50,100 and 200mM). (**C**) Absorbance values for different sugars (fructose, trehalose, galactose, D-glucose, sucralose and L-glucose) at 10mM concentration between high NaCl and normal media fed flies. (**D**) Mean feeding preference of wildtype (*CsBz*) flies for caffeine (10mM). In all graphs, black bars represent responses of flies pre-exposed to high NaCl diet compared to grey bars (normal fly media fed flies). (**E**) Mean feeding preference of wildtype (*CsBz*) flies at the indicated concentrations of other salts - sodium hydrogen carbonate (25 and 100mM), Di-sodium hydrogen O-phosphate (25 and 100mM), Magnesium Chloride (10mM and 100mM and potassium di hydrogen O-phosphate (10mM and 100mM). For absorbance assays in **A** and **C**, n= 2 sets each concentration, 60 flies each set. For feeding assays in **B**, **D** and **E**, N=6 plates, 20 flies each plate (10 males and 10 females). Statistical analysis was performed using ANOVA Tukey’s multiple comparison test for obtaining P values: *p < 0.05, **p < 0.005 and ***p < 0.0005. For all graphs, error bars=SEM and NS is not significant.

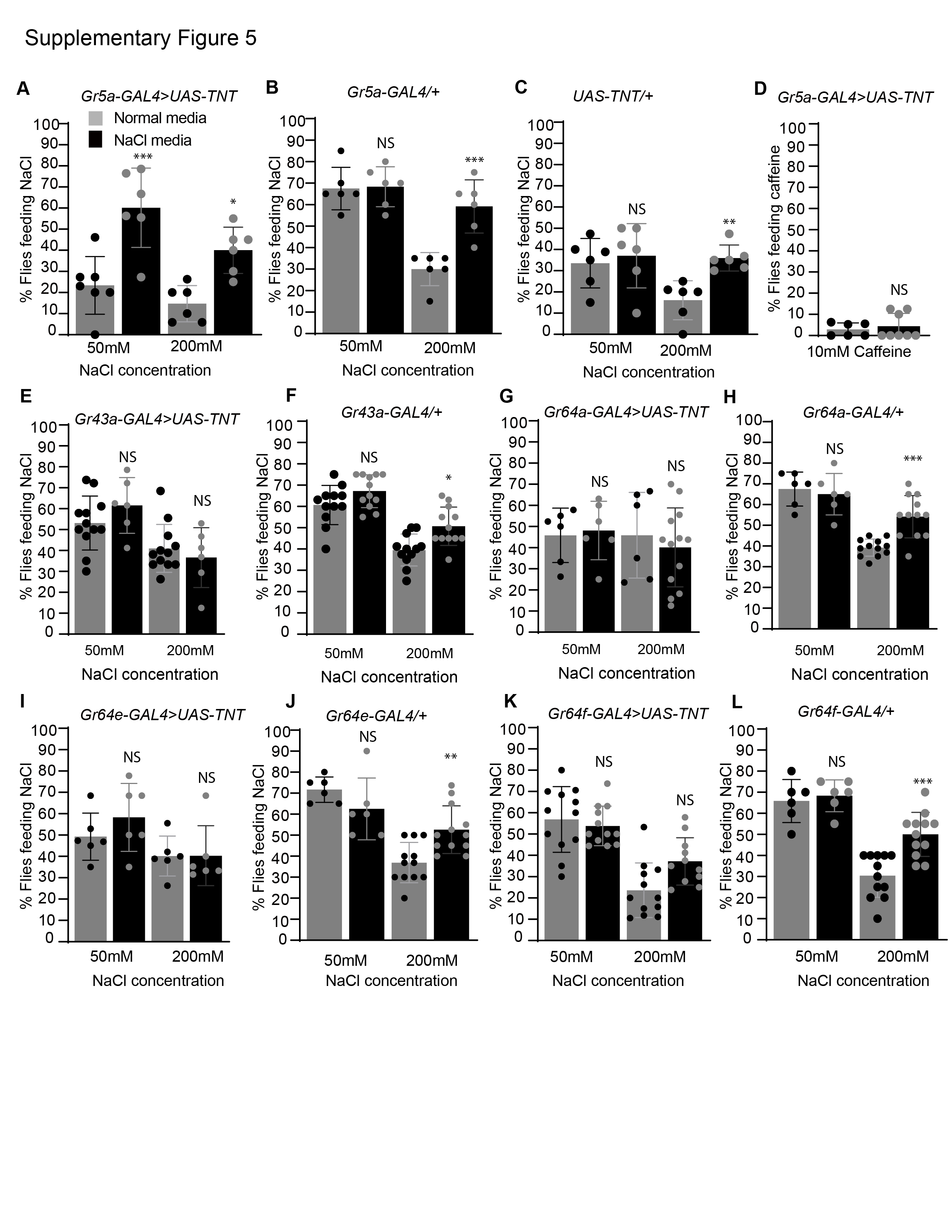

**SUPPLEMENTARY FIGURE 5**

**Role of neuronal activity in the peripheral and sweet LSO neurons of the pharynx in modulating high salt intake behavior under starvation**. (**A**) Mean feeding preference of *Gr5a-GAL4>UAS-TNT* flies after silencing neuronal activity of peripheral *Gr5a* sweet neurons by expressing tetanus toxin at low (50mM) and high (200 mM) NaCL concentrations after 24h of starvation. In all graphs, black bars represent responses of high NaCl-fed flies compared with normal media-fed flies (grey bars). Asterisks show significant differences between *Gr5a-GAL4>UAS-TNT* flies (black vs grey bars). (**B** and **C**) Mean feeding preference of parental flies (*Gr5a-GAL4/+* and *UAS-TNT/+).* (**D**) Mean feeding preference of *Gr5a-GAL4>UAS-TNT* flies (both normal media and salt pre-exposed) for caffeine (10mM). (**E**, **G**, **I**, and **K**) Mean feeding preference of flies after genetically manipulating the neuronal activity of other sweet taste receptor neurons by expressing tetanus toxin (*UAS-TNT*) using *Gr43a-GAL4*, *Gr64a-GAL4*, *Gr64e-GAL4*, and *Gr64f-GAL4* drivers, respectively. (**F**, **H**, **J,** and **L**) Mean feeding preference of parental flies (*Gr43a-GAL4/+, Gr64a-GAL4/+, Gr64e-GAL4/+* and *Gr64f-GAL4/+)* for 50 and 200 mM NaCl*.* For each graph, each bar, n=6-12 trails of 20 flies for each concentration (10 males and 10 females). Student’s t-test in was used for D otherwise statistical analysis was performed using ANOVA Tukey’s multiple comparison test for obtaining P values: *p < 0.05, **p < 0.005 and ***p < 0.0005. For all graphs, error bars=SEM and NS is not significant.

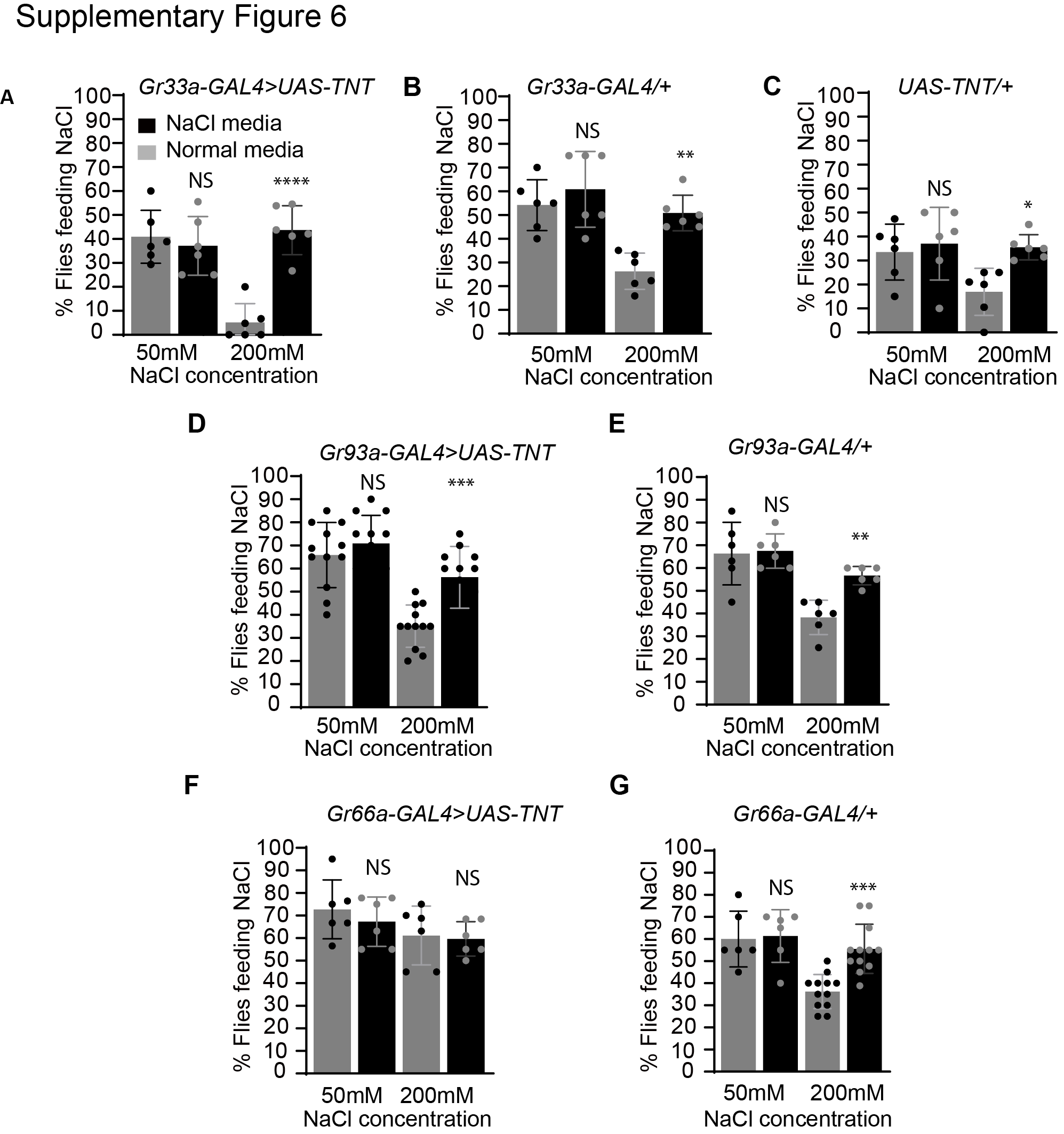

**Role of neuronal activity in the peripheral and bitter LSO neurons of the pharynx in modulating high salt intake behavior under starvation condition**. (**A**) Mean feeding preference of flies after genetically silencing activity of peripheral bitter neurons by expressing tetanus toxin (*Gr33a-GAL4>UAS-TNT*) at the indicated concentrations of NaCl after 24hrs of starvation. Asterisks show significant differences between *Gr33a-GAL4>UAS-TNT* flies fed on normal (grey bars) and salt media (black bars). (**B** and **C**) Mean feeding preference of parental control flies (*Gr33a-GAL4/+* and *UAS-TNT/+*). (**D** and **F**) Mean feeding preference of flies after silencing activity of other bitter neurons (*Gr93a* and *Gr66a*) by expressing *UAS-TNT* (**D**, *Gr93a-GAL4>UAS-TNT* and **F**, *Gr66a-GAL4>UAS-TNT*) between normal media and high NaCl media exposed flies (grey vs black bars). (**E** and **G**) Mean feeding preference of parental control flies (*Gr93a-GAL4/+* and *Gr66a-GAL4/+*). For each graph, each bar, n=6-12 trails of 20 flies for each concentration (10 males and 10 females). Student’s t-test in was used for D otherwise statistical analysis was performed using ANOVA Tukey’s multiple comparison test for obtaining P values: *p < 0.05, **p < 0.005 and ***p < 0.0005. For all graphs, error bars=SEM and NS is not significant.

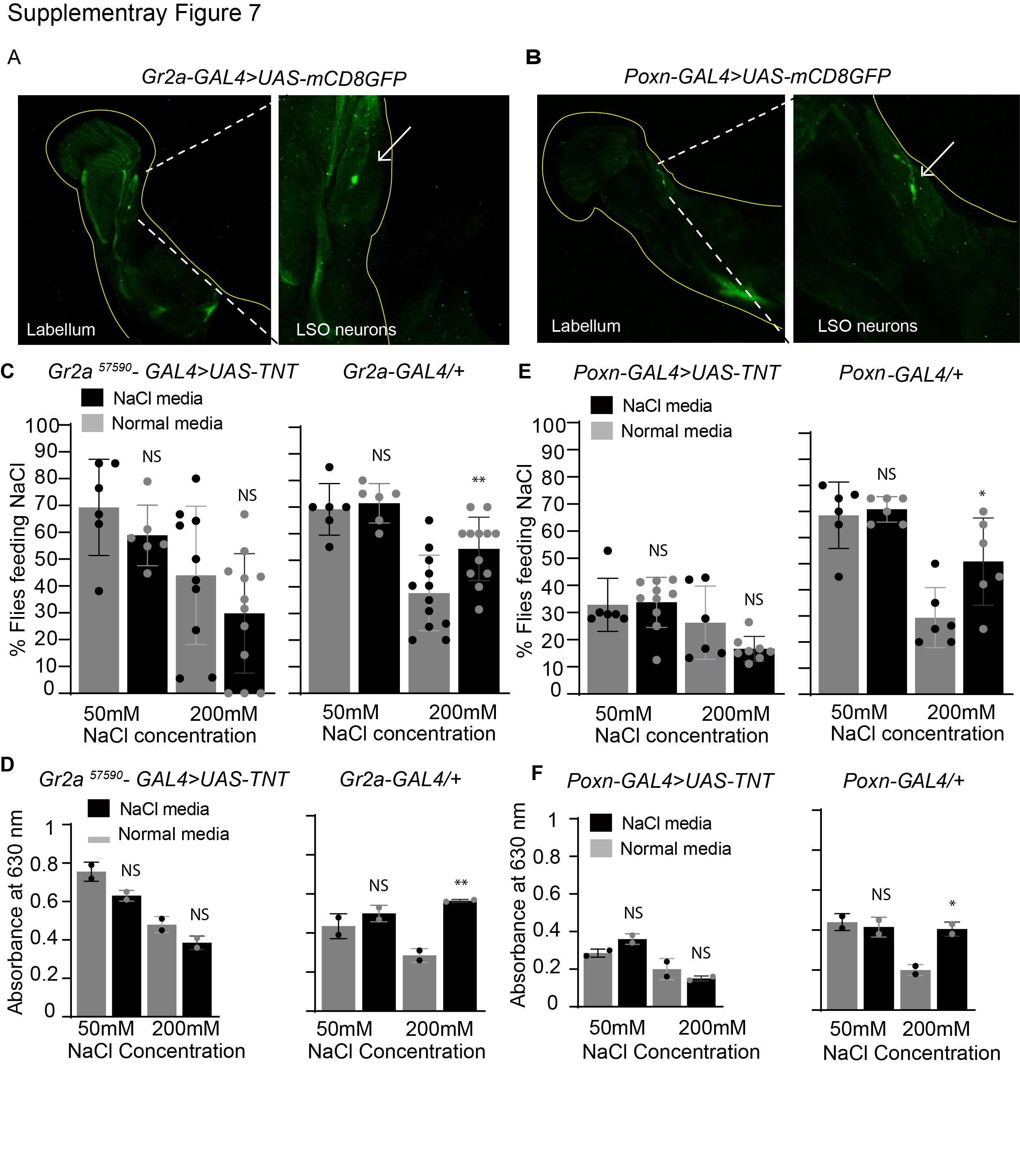

**SUPPLEMENTARY FIGURE 7**

**Silencing LSO pharyngeal neurons shows no increase in feeding preference for NaCl in high salt-fed flies.** (**A**) Expression pattern of *Gr2a-GAL4* labeled by *UAS-mCD8GFP* in the LSO pharyngeal neurons (white arrow in right panel). (**B**) Expression pattern of *poxn-GAL4* marked by *UAS-mCD8GFP* in the LSO pharyngeal neurons (white arrow in right panel). (**C**) Mean feeding preference of flies after silencing neuronal activity in the *Gr*2a (*Gr2a-GAL4>UAS-TNT* and parental control *Gr2a-GAL4/+*) positive LSO neurons tested with 50mM and 200mM NaCl (between normal media- and high salt media-fed flies; compare grey vs black bars). (**D**) Spectrophotometry analysis of normal and high salt media fed *Gr2a-GAL4>UAS-TNT* and *Gr2a-GAL4/+* flies. (**E**) Feeding preference of flies after silencing neuronal activity in the *poxn* (*poxn-GAL4>UAS-TNT* and parental control *poxn-GAL4/+*) positive LSO neurons (compare grey vs black) tested with 50mM and 200mM NaCl. (**F**) Spectrophotometry analysis of normal media and high salt media *poxn-GAL4>UAS-TNT* and *poxn-GAL4/+* flies. For **C** and **E**, n=6-12 plates, 20 flies each plate. For spectrophotometry analysis (**D** and **F**), n= 2 sets each concentration, 60 flies each set. Statistical analysis was performed using ANOVA Tukey’s multiple comparison test for obtaining P values: *p < 0.05, **p < 0.005 and ***p < 0.0005. For all graphs, error bars=SEM and NS is not significant.

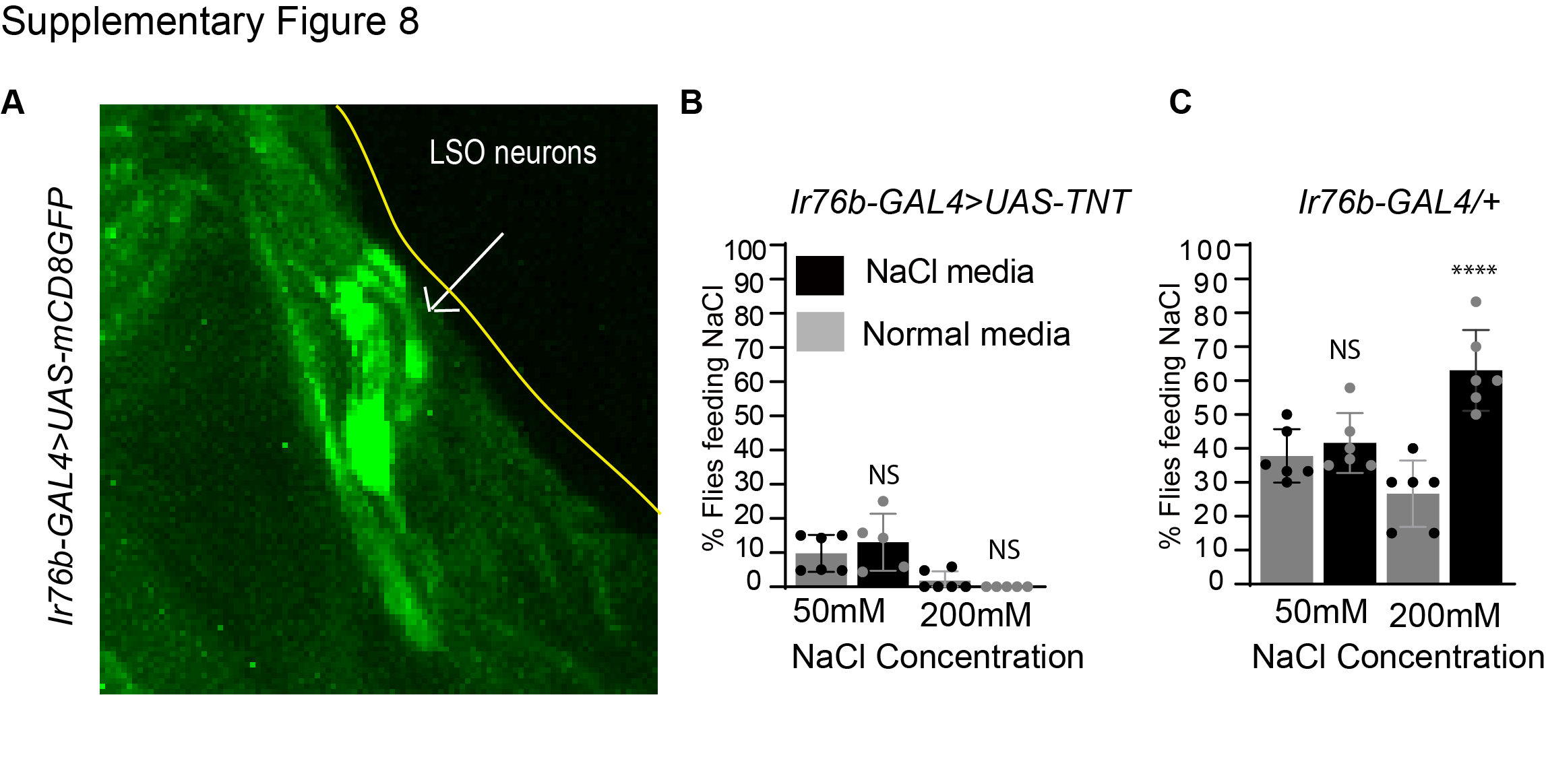

**SUPPLEMENTARY FIGURE 8**

**Silencing *Ir76b* LSO pharyngeal neurons showed no increased feeding preference for NaCl in high salt pre-exposed flies.** (**A**) Expression pattern of *Ir76b-GAL4* labeled with *UAS-mCD8GFP* in the LSO pharyngeal neurons (white arrow). (**B**) Mean feeding response of *Ir76b-GAL4>UAS-TNT* flies compared between normal media- and high salt media-fed flies. (**C**) Mean feeding preference of *Ir76b-GAL4/+* parental flies. Statistical analysis was performed using ANOVA Tukey’s multiple comparison test for obtaining P values: *p < 0.05, **p < 0.005 and ***p < 0.0005. For all graphs, error bars=SEM and NS is not significant.

**Supplementary movie 1 for Figure 2.**

Tarsal PER - Fly showing extension of proboscis when 50mM NaCl was presented

**Supplementary movie 2 for Figure 2.**

Tarsal PER - Fly showing no extension of proboscis when 200mM NaCl was presented.
